# The control of epistemic curiosity in the human brain

**DOI:** 10.1101/157644

**Authors:** Romain Ligneul, Martial Mermillod, Tiffany Morisseau

## Abstract

Epistemic curiosity (EC) is a cornerstone of human cognition that contributes to the actualization of our cognitive potential by stimulating a myriad of information-seeking behaviours. Yet, its fundamental relationship with uncertainty remains poorly understood, which limits our ability to predict within- and between-individual variability in the willingness to acquire knowledge. Here, a two-step stochastic trivia quiz designed to induce curiosity and manipulate answer uncertainty provided behavioural and neural evidence for an integrative model of EC inspired from predictive coding. More precisely, our behavioural data indicated an inverse relationship between average surprise and EC levels, which depended upon hemodynamic activity in the rostrolateral prefrontal cortex from one trial to another and from one individual to another. Complementary, the elicitation of epistemic surprise and the relief of acute curiosity states were respectively related to ventromedial prefrontal cortex and ventral striatum activity. Taken together, our results account for the temporal evolution of EC over time, as well as for the interplay of EC, prior knowledge and surprise in controlling memory gain.

## Introduction

Epistemic curiosity (EC) predicts educational success (Von Stumm et al., 2011), orients our attention (Gottlieb et al., 2013; Kidd and Hayden, 2015) and underlies many decisions of our everyday life, such as opening books, browsing the internet, watching movies or engaging in trivia quizzes. Following Aristotle’s thesis that “all men by nature desire to know”, it has long been suggested that knowledge might act as an intrinsic reward and curiosity as an innate drive in humans. Largely embraced by previous neuroimaging studies of EC (Gruber et al., 2014; Kang et al., 2009), this view may explain why EC seems to outreach the information required to optimize survival and reproduction, as it extends to cultural domains with unclear or indirect biological value such as philosophy, science and art. However, by turning knowledge itself into a primary biological goal, one leaves aside most of the developmental and situational determinants of curiosity responsible for the large inter- and intra-individual variability associated with information-seeking behaviors. Is it possible to explain the extent and the regulation of human curiosity without relying exclusively on the hypothesis that curiosity depends an innate “thirst of knowledge” susceptible to satiation? Recent theoretical developments have indeed opened the path to a more complete, yet complex, picture which still awaits empirical validation.

EC is typically triggered by the awareness of specific gaps in one’s own knowledge — hence directing exploration towards the information expected to fill these gaps (Berlyne, 1966; Gottlieb et al., 2013; Loewenstein, 1994). Yet, psychology research has shown this process varies greatly across individuals and contexts. For example, some individuals may actively avoid an information (Gigerenzer and Garcia-Retamero, 2017; Sweeny et al., 2010) while others may seek it despite enduring risks (Hsee and Ruan, 2016; Pierce et al., 2005; van Dijk and Zeelenberg, 2007). Tolerance to uncertainty has recently emerged as a key factor underlying this variability (Kashdan et al., 2009; Litman, 2010), as the outcome of exploration does not always fill the knowledge gaps under scrutiny but can also expose individuals to unexpected information. In turn, exposure to unexpected information may challenge the sense of coherence or completeness in adjacent knowledge domains, eventually producing a net increase in perceived ignorance or uncertainty. For example, if a neuroscientist attempting to confirm the involvement of brain region A in a given cognitive process eventually observes that brain region B co-activates with A, the unexpected information carried by the activation of B may dampen his or her confidence regarding the experiment as a whole. Following a typology early proposed by Berlyne (Berlyne, 1966, 1954), this distinction between the willingness to acquire the specific information missing to solve existing problems (often termed “specific” or “deprivation” EC) and the willingness to process new information in general (often termed “nonspecific” or “diversive” EC) is at the heart of the current study. Indeed, while these two facets of EC are known to differ at the personality level, both in the general population (Litman, 2010; Litman and Spielberger, 2003) and in academia (where it predicts deeper *versus* broader contributions to scientific progress (Bateman and Hess, 2015), their cognitive and neural underpinnings remain largely obscure.

Predictive coding constitutes a promising framework to understand how the specific and nonspecific components of EC are triggered and regulated by context or reinforced over time. The core principle of predictive coding theories is that the reduction of uncertainty and surprise constitutes the primary function of our cognitive systems (Clark, 2013; Friston et al., 2012). In this framework, it is thus natural for acute uncertainty and ignorance states to be associated with negative affects requiring relief (Hirsh et al., 2012; Litman et al., 2005; Loewenstein, 1994). However, this powerful principle must be reconciled with the manifold curiosity-related behaviours which can transiently increase uncertainty (e.g pressing the “random button” of Wikipedia) and even lead to sustained doubtful states (e.g reading Descartes). Indeed, avoiding further stimulation and refraining from acting often appears an efficient way to escape new sources of uncertainty, once existing ones have been addressed (Clark, 2013; Friston et al., 2012). This objection known as the “dark room” problem argues against the existence of nonspecific EC, which orients us towards new sources of uncertainty. Proponents of predictive coding have thus proposed that individuals might be born with (or develop) second-order expectations regarding the optimal amount of uncertainty which should be experienced when interacting with their environment (Clark, 2013; Friston et al., 2015; Schwartenbeck et al., 2013). Inspired by early optimal arousal theories (Berlyne, 1966; Hebb, 1955), this second principle implies that humans may be “surprised of not being surprised” and that they may actively try to fulfill their surprise expectations by engaging exploration, whenever their uncertainty levels are lower than expected (and vice-versa). Therefore, transient surprise signals may not only modulate confidence in one’s own knowledge or increase memory gain, they may also update an estimate of the average surprise experienced by the organism so that nonspecific EC levels can be adjusted upwards or downwards in order to match this estimate with expectations, through a mechanism sometimes called “active inference” (Friston et al., 2015). Importantly, such conceptualization suggests that distinct psychological processes and neural circuits may underlie each EC dimension.

On the one hand, the nonspecific motivation to obtain information and to confront new epistemic problems should depend on a neural representation of uncertainty dependent upon average surprise. Recently, the combination of computational modelling and neuroimaging techniques showed that neural activity in the rostrolateral cortex (rlPFC) mediates the control of exploration by uncertainty in foraging tasks (Badre et al., 2012; Boorman et al., 2009; Donoso et al., 2014). As exogenous electrical inhibition of the rlPFC causally reduces exploration in such contexts(Raja Beharelle et al., 2015), rlPFC activity may underpin the putative relationship between nonspecific EC and average surprise, as expected from the predictive coding framework. Yet, another network is likely required to compute surprise signals in the first place, as this process entails a complex comparison of incoming information with prior knowledge. Together with the hippocampus, the ventromedial prefrontal cortex (vmPFC) appears as a good candidate to perform such task, as recent studies highlighted its role for online memory retrieval(Lighthall et al., 2014; Rissman et al., 2016), for schema-based memory (Garrido et al., 2015; Ghosh et al., 2014; van Kesteren et al., 2013) and for representing confidence in one’s own knowledge (Lebreton et al., 2015). On the other hand, the motivation to reduce specific sources of uncertainty may depend on a “reinforcement by relief” (of ignorance) recruiting the reward system, likewise pain relief (Navratilova and Porreca, 2014; Seymour et al., 2005). In line with this hypothesis, Jepma and colleagues recently demonstrated that relieving perceptual curiosity elicits higher BOLD responses in the ventral striatum (Jepma et al., 2012), a key subcortical region involved in numerous reinforcement-learning paradigm (Garrison et al., 2013). Yet, the involvement of these neurocognitive processes in epistemic curiosity *per se* remains to be demonstrated.

In the current study, we used a two-step stochastic trivia quiz (Fig. 1a) designed to induce and manipulate curiosity levels in 22 participants undergoing fMRI (see SI Materials and Methods for extensive details about the paradigm). Our quiz focused on cinema because of the widespread interest in this domain across sexes, cultures, and education levels. This choice also facilitated the standardization of answers (only movie titles) and the evaluation of prior knowledge related to answers (titles were categorized as watched or not by the participant). During the first part of the quiz (run 1), participants rated their curiosity for 60 cinema-related questions. After each rating, the answer to the question was either revealed (50%) or replaced by hashtags (50%) to manipulate answer uncertainty and maximize variations in the nonspecific EC component. In the second part (run 2), the same 60 questions were presented again and participants were asked to indicate whether they could spontaneously remember the answers. At this point, questions that had not been answered in run 1 could still elicit curiosity and their associated (not yet revealed) answers could still elicit surprise, whereas remembered items served as controls, matched with the former in terms of visual stimulation and linguistic content. After the main task, a localizer involving individualized sets of new movie titles was used to reveal the brain regions responding to prior knowledge in a task-independent fashion (run 3). Unannounced post-test questionnaires were finally administered outside the scanner, including a recall test as well as surprise and interest ratings for each item.

**Figure 1.**
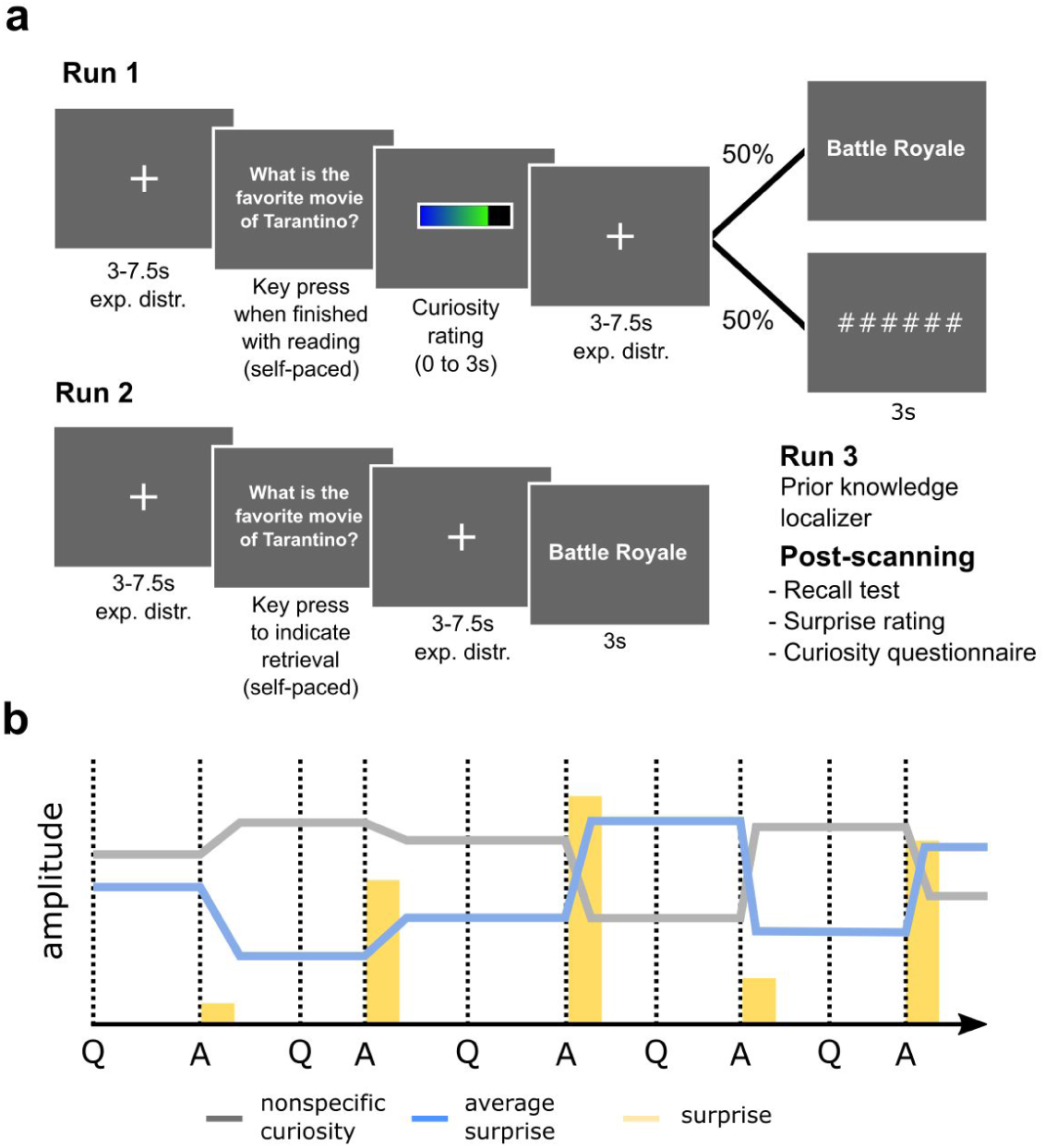
Design of the fMRI study task and main hypothesis. (a) In run 1, participants were presented with 60 prescreened trivia items (see also Table S1). After reporting curiosity on a non-numerical continuous gauge, they were presented with either the answer (for half of the trivias), or hashtags (for the other half). In run 2, participants were presented again with the 60 questions and reported whether the answer came spontaneously to their mind or not. Each answer was then revealed, so that half of the answers relieved curiosity whereas the other half merely echoed a previously encountered information. In run 3, participants were presented with an independent set of movie titles they had watched or not (prior knowledge localizer). Once outside the MRI scanner, they were finally asked to report all the answers they could remember and to rate their surprise and interest levels for each trivia answer. (b) Schematic representation of the opponency expected from the predictive coding framework between nonspecific EC (in grey) and the average amount of surprise experienced (in blue) during a series of questions (Q) and answers (A) eliciting different amounts of epistemic surprise (in yellow).

## Results

### The interplay of prior knowledge, curiosity and surprise for memory encoding

Behavioural analyses demonstrated that possessing some prior knowledge related to the answers (as assessed by whether or not the participant had seen the target movie before the experiment) increased both curiosity ratings (t(21)=2.46, p=0.023) and surprise ratings (t(21)=4.86, p<0.001) (Fig. 2a). Curiosity and surprise were positively correlated to each other (r=0.22, p<0.001, ratings z-scored for each participant individually; Fig. 2b) and a repeated-measures ANOVA confirmed that curiosity predicted surprise ratings (F(2,42)=27.4, p<0.001). Moreover, curiosity, surprise and prior knowledge were all positively associated with recall performances (whether the response was correct or not in the post-scan memory test) according to median-split analyses separating items as a function of high and low curiosity (z=3.88, p<0.001), surprise (z=3.32, p<0.001) and prior knowledge (z=3.81, p<0.001; Fig. 2c).

**Figure 2.**
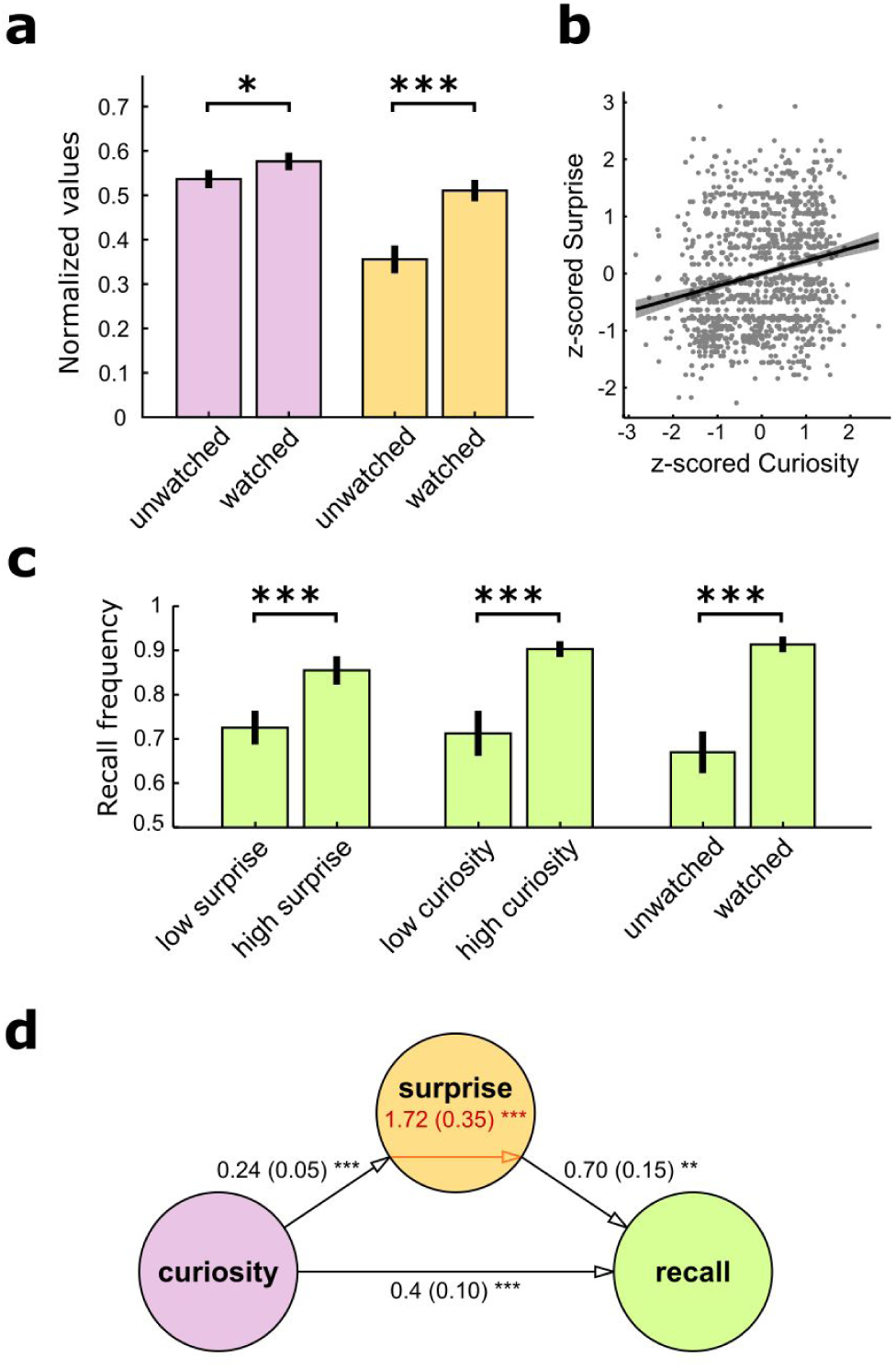
Behavioural results. (a) The presence of prior knowledge about target answers (whether or not the participant had seen the movie before the experiment) was associated with increased curiosity ratings (rose) and surprise ratings (yellow) in post-scan questionnaires (ratings rescaled between 0 and 1). (b) Higher curiosity levels were also associated with increased surprise ratings. (c) Recall performances were affected by surprise, curiosity and prior knowledge, as revealed by median-split analyses. (d) Participant-wise mediation analyses demonstrated that curiosity induced direct and surprise-mediated benefits for the ability to recall answers in the post-test task (logistic mediation with unsuccessful recall coded as 0 and successful recall coded as 1, with prior knowledge and repetition included as covariates). Error bars represent s.e.m. p<0.05*, p<0.01**, p<0.001**.

In order to exclude the possibility that prior knowledge, curiosity and surprise would simply reflect a common latent variable (e.g. attention, affective value), we used a Generalized Estimating Equations approach (see SI Material and Methods). The three factors were individually significant (curiosity: beta=1.27, Wald χ2 = 15.9, p<0.001; surprise: beta=0.43, Wald χ2 = 10.2, p=0.001; prior knowledge: beta = 1.58, Wald χ2 = 74.4, p<0.001), hence confirming their complementary contribution to memory encoding.

A logistic mediation analysis including prior knowledge and condition (whether the answer had been seen once or twice during the experiment) as covariates further showed that surprise partly mediated the beneficial effects of curiosity on recall performances (indirect path: z=3.47, p<0.001; direct path: z=3.64, p<0.001; Fig. 2d). Finally, as expected from a previous study(McGillivray et al., 2015), items rated as more interesting were associated with higher surprise ratings (t(21)=2.48, p=0.02) but, contrary to surprise, interest did not mediate curiosity-driven memory benefits (indirect path: z=1.31, p=0.21). Therefore, despite its tight relationship with curiosity and prior knowledge, epistemic surprise reflects a specific construct that mediates curiosity-driven memory encoding.

### The neural correlates of epistemic curiosity

Before testing more specific hypotheses regarding its dynamical regulation by surprise, we investigated the neural correlates of EC ratings, in line with previous studies (Gruber et al., 2014; Kang et al., 2009). When analyzing curiosity-dependent BOLD responses at the answer stage (for which our paradigm was optimized), we observed a clear-cut encoding of curiosity relief in the ventral striatum during run 1 where curiosity was relieved in 50% of the trials (whole-brain analysis, voxel-wise threshold: p<0.005, cluster-wise threshold: p_FWE_<0.05; Fig. 3a). This conclusion held when the analysis was restricted to an anatomical mask (Ahsan et al., 2007) of the nucleus accumbens (NAcc ROI: t(21)=2.54, p=0.02), while no significant modulation was observed when hashtags were displayed (NAcc ROI: t=0.03, p=0.97). Yet, in run 2 (where curiosity was relieved in 100% of the trials), no modulation by EC was observed when participants processed the answers which had not been delivered in run 1 (p>0.2; Fig. 3b).

**Figure 3.**
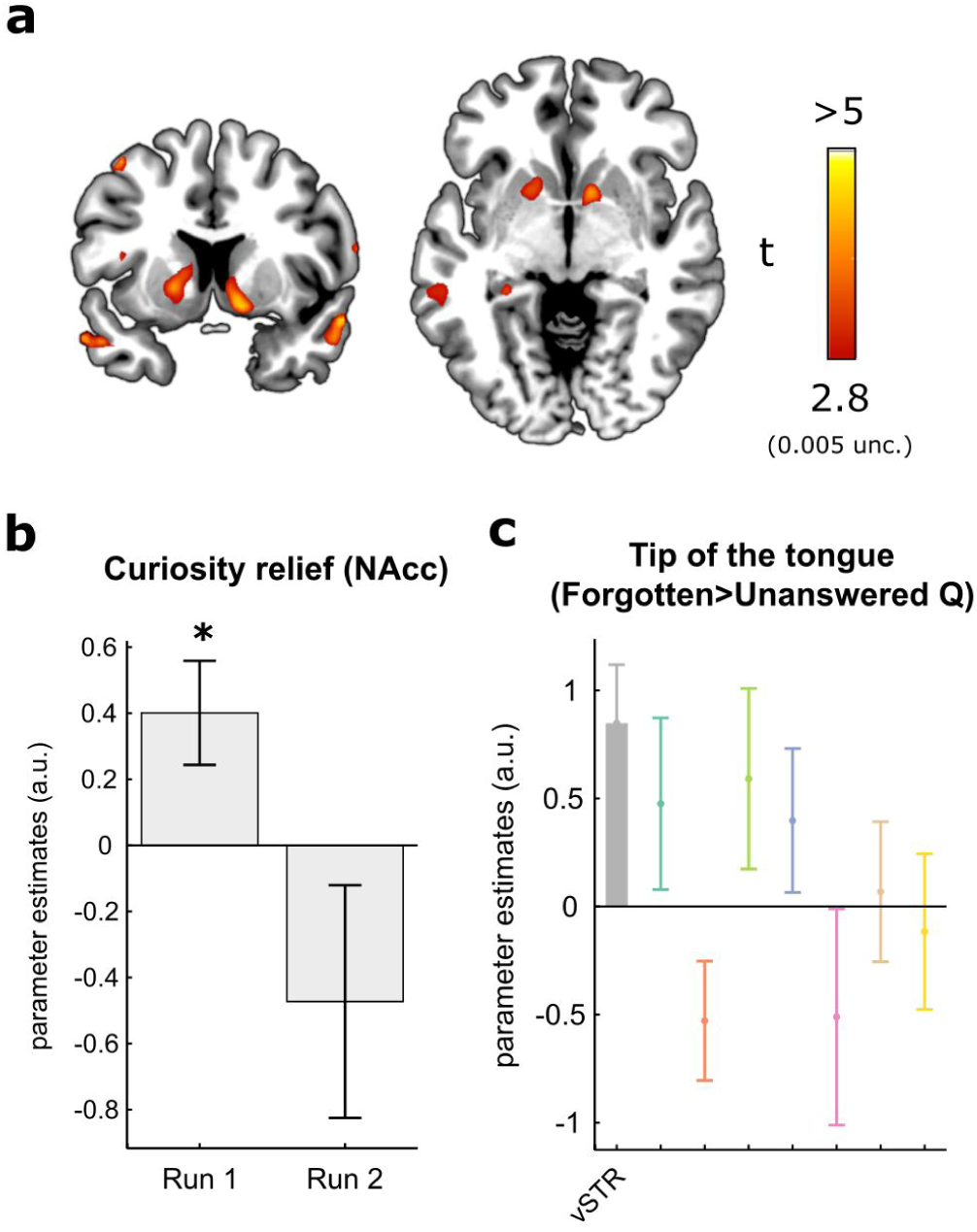
Neural activities related to epistemic curiosity in the ventral striatum. (a) Curiosity levels modulated ventral striatal responses to knowledge delivery during the stochastic trivia quiz (voxel-wise threshold: p=0.005^UNC^; cluster-wise threshold: p<0.05^FWE^; MNI peak: [−6 5 1]). (b) In the first run, the effect reported in **a** was also significant in the nucleus accumbens (NAcc) but it disappeared during the second part of the trivia quiz. **c**, Among the 8 ROIs described in Fig. 5b (same color code), the ventral striatum was the only region to activate significantly more in response to old but forgotten questions in run 2, as compared to never answered questions. Error bars represent s.e.m. p<0.05* (two-tailed).

At the question stage, whole-brain analyses showed that more intense curiosity states induced more activity in the dmPFC in run 1 (Fig. S1a) but not in run 2 (Fig. S1b). No other cluster reached the standard threshold for multiple comparisons in either run (p_FWE_<0.05). Although we did not observe any striatal encoding of EC as reported in(Gruber et al., 2014), it must be stressed that our task was not optimized to investigate BOLD responses at the question stage (e.g variable question length, presence of motor responses). Clear-cut activations were observed in the striatum alongside other areas such as the hippocampus when participants processed questions for which they knew answers as compared to questions for which answers were unknown, in both runs (Fig. S1c-d; NAcc ROI, run 1: z(15)=3.25, p=0.001; excluding 6 participants who ignored all answers; NAcc ROI, run 2: t(21)=3.07, p=0.006). Interestingly, the ventral striatum was the only region activating more in front of questions previously answered in run 1 but forgotten in run 2, as compared to never answered questions (t(21)=3.13, p=0.005; Fig. 3c). This pattern suggests that specific memory retrieval and motivational processes were somewhat active in the second run of our task.

### Surprise-dependent control of curiosity

Our main hypothesis concerned the variation of nonspecific curiosity levels over time, as a function of the average amount of surprise recently experienced (Fig. 1b). In the computational approach used to tackle this issue (SI Materials and Methods), we assumed that the subjective levels of curiosity reported in participants’ ratings during run 1 resulted from two distinct additive influences: (i) the motivation to relieve an acute ignorance state induced by the specific question displayed on screen (specific EC); (ii) the motivation to be exposed to any new information (nonspecific EC) conceived as an item-independent variable fluctuating slowly throughout the quiz. Importantly, the delta-rule algorithm used to monitor the average amount of surprise was totally blind to the content of the questions and to the outcome of a trial *t*: consequently, it could only explain the variance associated with this nonspecific component of EC based on previous items (*t-1, t-2*, etc).

Supporting our hypothesis, a delta-rule that monitored average surprise in each trial (Q{sur}) outperformed the model that monitored only the frequency of knowledge delivery (Q{0-1}) and models that included time as a regressor, either alone or in combination with any of the two delta-rules (Fig. 4A). Bayesian group comparisons treating model attribution as a random effect indicated that this conclusion held both when curiosity ratings were considered as a normally distributed variable and when they were binarized into high/low categories and predicted by means of logistic regression (see Methods and Supplementary Fig. S2a-e for extensive details). Crucially, the overall effect of expected surprise on curiosity ratings was negative in both the continuous (t(21)=-2.95, p=0.008) and the binomial cases (t(21)=-3, p=0.007), which confirmed that low average surprise exacerbated nonspecific EC whereas high average surprise suppressed it — as expected from the predictive coding logics.

**Figure 4.**
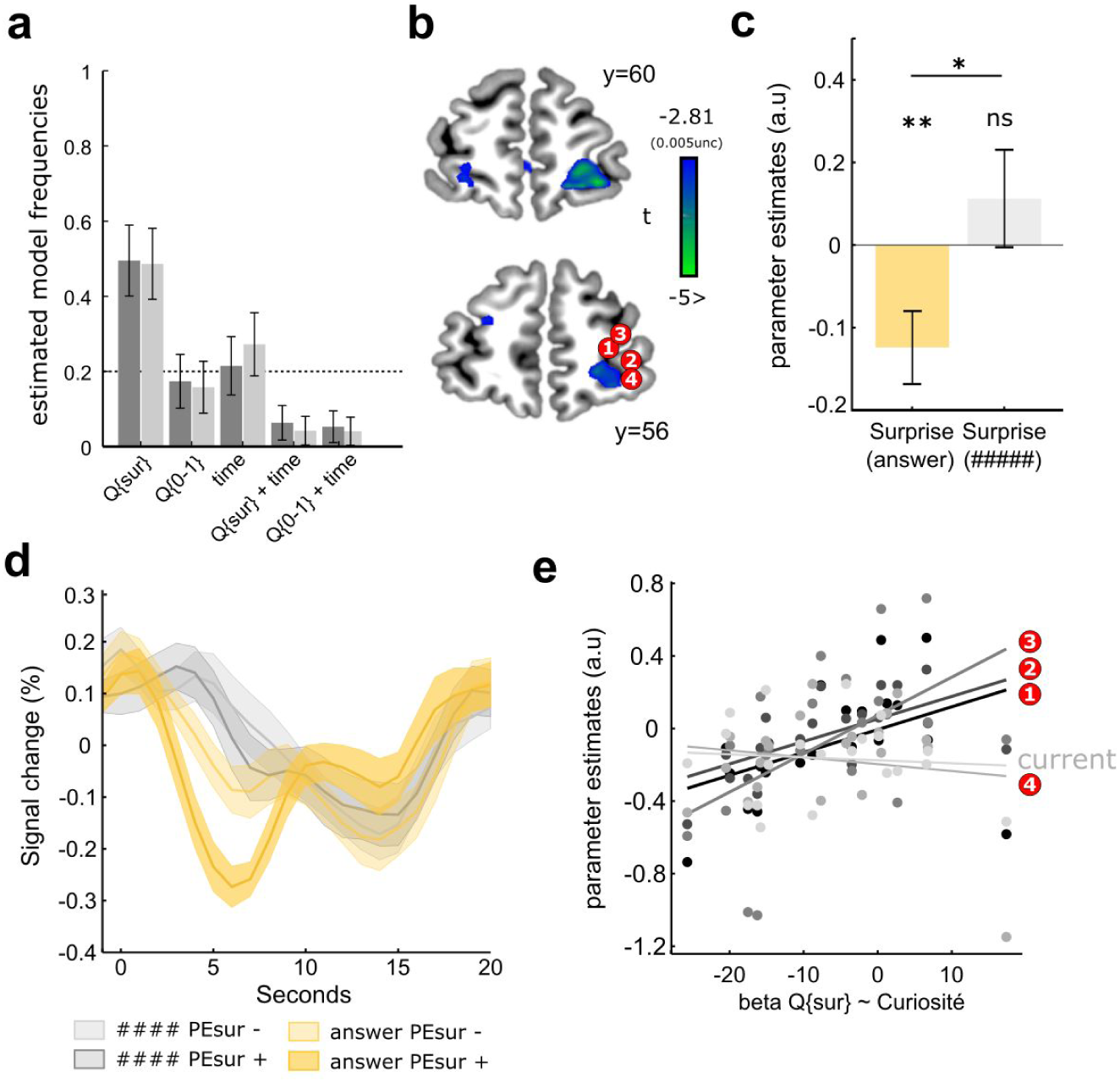
Surprise-dependent control of curiosity and the rlPFC (see also Supp. Fig. S2). (a) The model that updated on each trial the average amount of surprise recently experienced outperformed the four alternative models tested to account for the evolution of curiosity ratings over time, for both the Akaike (dark grey) and Bayesian (light grey) Information Criteria. (b) Neural correlates of surprise prediction errors at the answer stage (run 1) in the right rostrolateral prefrontal cortex. This functional cluster was close but did not overlap the activation peaks reported in the literature on uncertainty-driven exploration. (c) The rlPFC also correlated with surprise ratings themselves when answers but not hashtag were delivered (3mm sphere around peak reported in 4b). (d) Finite Impulse Response modelling confirmed the presence of genuine rlPFC deactivations in responses to stronger surprise prediction errors, occuring only when answers were actually delivered (yellow).The encoding of PE{sur} at peaks 1-3 was correlated with the the suppression of nonspecific curiosity by Q{sur}, as observed at the behavioural level. p<0.05*, p<0.01**, p<0.001*** (two-tailed). Error bars represent s.e.m. Plotted signals were extracted from 3mm-radius spheres centered around the peaks of interest.

### Monitoring of average surprise and regulation of curiosity by the rostrolateral PFC

Next, we studied how the brain tracked changes in average surprise from one trial to another. To do so, we investigated the parametric encoding of surprise prediction errors (PE{sur}) at the answer stage of the run 1, during which variations in EC were assessed. Formally, this trial-wise variable corresponds to the surprise experienced in each trial minus the average surprise recently experienced. Restricted to a prefrontal mask spanning all voxels anterior to the head of the caudate (Y>22), our analyses revealed a significantly higher level of hemodynamic activity related to PE{sur} within the right rostrolateral prefrontal cortex (p<0.05^FWE^, MNI [18 62 −11], Fig. 4b). Importantly, the GLM excluded potential confounding effects of displaying answers *versus* hashtags (modeled as separate events), as well as those associated curiosity relief and prior knowledge related to answers (both included before PE{sur} in the serial orthogonalization procedure implemented by SPM, see Methods). Additionally, replacing PE{sur} by surprise ratings in the GLM confirmed that the rlPFC not only encoded surprise prediction errors but also surprise itself (t(21)=-2.85, p=0.009; Fig. 4c). Finally, since parameter estimates were negative, we used a Finite Impulse Response model distinguishing outcomes (answer and hashtag) as a function of PE{sur} to ascertain that stronger surprise prediction errors triggered proportional deactivations of the rlPFC (subject-wise median-split; Fig. 4d).

Given that the cluster reported in Fig. 4b appeared more anterior and medial than expected based on the literature on uncertainty-driven exploration, we also assessed the effect of PE{sur} at 4 previously reported rlPFC coordinates (Badre et al., 2012; Boorman et al., 2009; Daw et al., 2006; Donoso et al., 2014). No significant modulations were observed for PE{sur} (all p>0.2), but the encoding of PE{sur} predicted the influence of this uncertainty-related variable on curiosity ratings from one subject to another at 3 peaks out of 4 (Fig. 4e; peak from Daw et al: ρ=0.61, p=0.003; Boorman et al: ρ=0.69, p<0.001; Donoso et al., ρ=0.53, p=0.014) while it was not the case at the peak reported in Badre et al. (ρ=0.24, p=0.29) or at the peak reported Fig. 4B (ρ=-0.09, p=0.68). In other words, a more posterior and lateral portion of the rlPFC seemed to implement the control of nonspecific EC levels based on computations performed in the more medial and anterior site highlighted by our initial analysis.

### Genesis of epistemic surprise in the ventromedial PFC

Having established that the rlPFC monitors an average surprise variable and implement its influence on EC, we sought to characterize the neural origin of the epistemic surprise signal itself. To do so, we reasoned that the area computing epistemic surprise should be sensitive: (i) to information delivery, (ii) to information novelty, (iii) to prior knowledge, and (iv) to surprise itself. This selectivity profile followed from the definition of epistemic surprise as the mismatch between new information and prior knowledge. Since our experiment did not provide the statistical power required for a 4-way conjunction analysis, we chose to investigate the selectivity profile of all areas which survived multiple comparisons (p<0.05^FWE^) for the contrast “answer > hashtag” (Fig. 5a) and for the parametric effects of EC at the question and answer stages, also obtained from the first run (for details see Methods). Clusters in the dorsolateral prefrontal cortex (dlPFC), the hippocampus (HPC), the precuneus (PrC), the superior temporal sulcus (STS), the ventromedial PFC (vmPFC), the dorsomedial PFC (dmPFC) and the ventral striatum (vStr) revealed by these contrasts were thus transformed in bilateral regions of interest (ROIs) for the following analyses (Fig. 5b).

**Figure 5.**
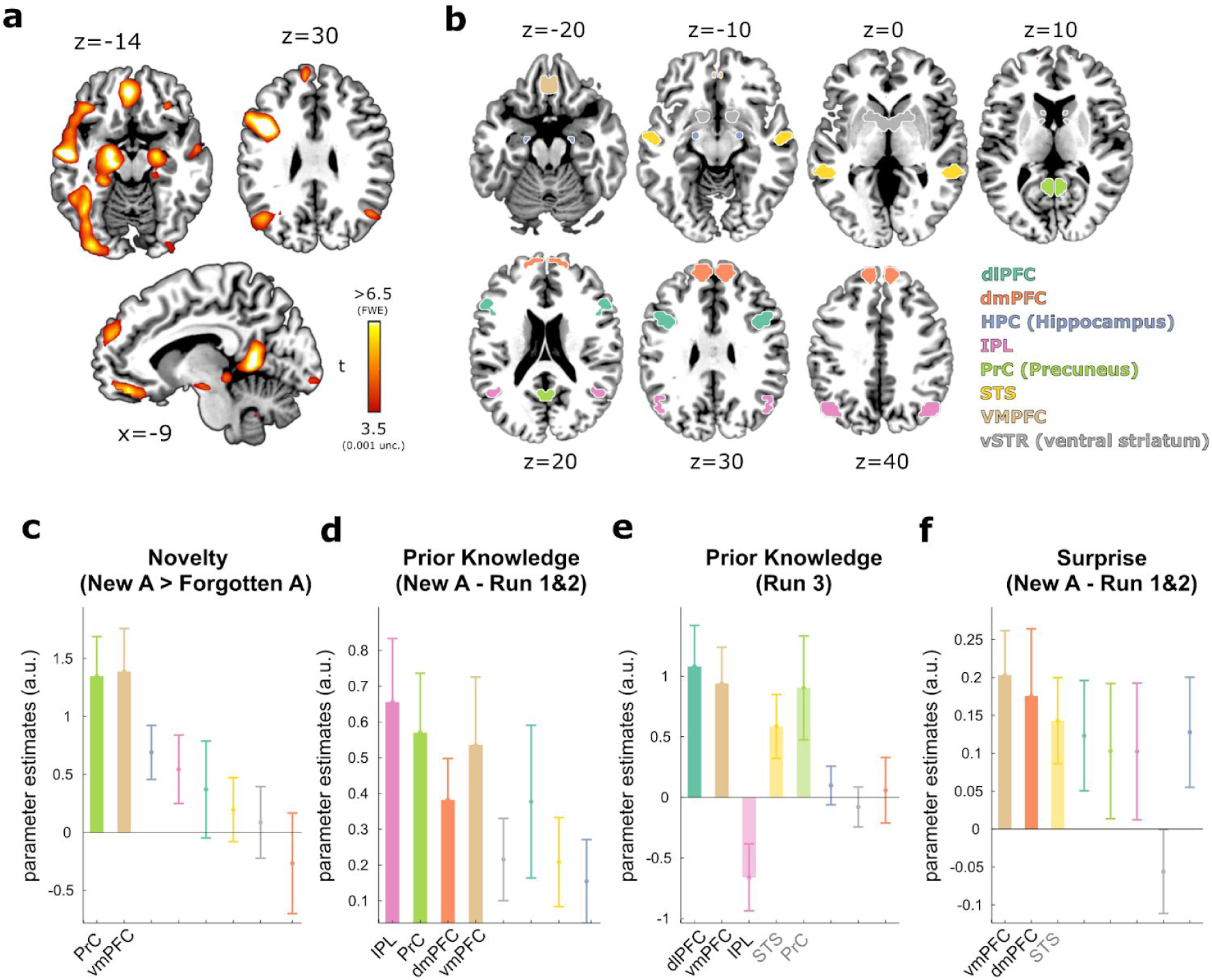
Epistemic surprise signals deriving from the comparison of new information with prior knowledge in the vmPFC (see also Fig. S1). (a) Among other areas, the vmPFC was more activated when processing new answers than hashtags (run 1, p<0.05^FWE^, MNI peak: [−3, 35, −17]) or old answers (run 2, see Table S4). (b) Eight bilateral ROIs were systematically investigated to highlight the central role of the vmPFC in the genesis of epistemic surprise (see also Fig. S2a and Fig. 6a). (c) Sensitivity to information novelty was revealed by comparing new answers to old but forgotten answers in the second run. (d) Encoding of prior knowledge pooled over the two runs of the trivia quiz when processing new answers (watched *versus* unwatched movie titles). (e) Encoding of prior knowledge in the localizer task which presented participants with a separate set of watched or unwatched movie titles, independently of any trivia question (run 3). (f) Encoding of surprise ratings associated with new answers, pooled over the two runs of the trivia quiz. In graphs c-f, areas surviving the p(FDR)<0.05 are plotted with plain colors, areas significant only at an uncorrected threshold (p<0.05) are plotted with half-transparent colors and non-significant effects are reported using only error bars. Effect are ordered from left to right as a function of their significance. Error bars represent s.e.m.

Confirming their general sensitivity to knowledge delivery, the 5 areas which responded to the contrast “answer > hashtag” in run 1 also activated significantly more in response to new answers as compared to old and remembered answers in run 2 (Fig. S1e). The precuneus (t(21)=3.94, p<0.001) and the vmPFC (t(21)=3.74, p=0.001) were the only regions discriminating new answers from old and forgotten answers, hence implying a genuine sensitivity to information novelty (Fig. 5c). Moreover, the vmPFC was the only region to encode prior knowledge about answers both in the main task (t(21)=2.82, p=0.010; Fig. 5d) and during the localizer task (t(21)=3.14, p=0.005; Fig. 5e), although less pronounced contributions of the precuneus and the dlPFC were also observed. Crucially, when new answers were delivered (run 1 and 2), the vmPFC activated more in response to higher epistemic surprise (t(21)=3.47, p=0.002) alongside the STS (t(21)=2.52, p=0.019; Fig. 5f). Excluding that this latter effect reflected a broader “valuation” process, vmPFC activity did not reflect surprise ratings when participants viewed old answers in run 2 (t(21)=-1.70, p=0.10, Fig. S1f; difference new *versus* old: t(21)=2.65, p=0.011, paired t-test) nor interest ratings (run 1 and 2: t(21)=1.18, p=0.25; Fig. S1g). Note also that the surprise regressor was orthogonalized on curiosity and prior knowledge, hence excluding that the observed effect would reflect its correlation with these variables.

Consistent with the interplay of prior knowledge, surprise and recall observed at the behavioural level, the vmPFC was the only region whose activation at the time of new answers predicted subsequent recall in the post-scan memory test (t(21)=2.21, p=0.03; Fig. S1h). Although this effect did not survive the false-discovery rate criterion used to correct for multiple comparisons across ROIs, it remains remarkable as the regressor indexing “subsequent recall” was orthogonalized on curiosity, prior knowledge and PE{sur}.

## Discussion

Our study provides evidence for a key assumption of the predictive coding framework regarding the evolution of epistemic curiosity (EC) over time. Indeed, when the average surprise elicited by previous trivia answers was high, upcoming EC ratings were low, and *vice-versa*. Neuroimaging data revealed that this regulatory mechanism was implemented by the rostrolateral prefrontal cortex (rlPFC), whose activity encoded surprise prediction errors when processing trivia answers. Facilitating memory encoding, surprise itself was presumably computed within the ventromedial prefrontal cortex (vmPFC) in relationship with other brain areas sensitive to prior knowledge and information novelty. Moreover, we show that the relief of acute EC states triggered by trivia questions recruits the reward system when knowledge delivery is stochastic, a condition which was not met in earlier neuroimaging studies of EC. Taken together, these results shed light on the neural circuitry underlying the specific and nonspecific components of EC and suggest that disentangling these two dimensions will be crucial to better understand the origins of within- and between-subject variability in this fundamental cognitive trait.

Previous neuroimaging studies of epistemic curiosity showed that reading questions eliciting higher curiosity produced higher BOLD responses of the striatum, a key player of the so-called brain reward system (Gruber et al., 2014; Kang et al., 2009). Based on these findings, epistemic curiosity was recently reframed as a reward-seeking process where “reward is information” (Marvin and Shohamy, 2016). Although this view may be attractive for its simplicity and certainly captures some features of EC, the hypotheses tested in the current study were instead rooted in the “predictive coding” framework which is also able to account for the dependence of curiosity upon prior knowledge and surprise. In this framework, EC primarily depends on uncertainty-minimization rather than reward-maximization, which means that higher curiosity ratings do not primarily reflect the anticipation of more pleasurable outcomes but the experience of more intense states of uncertainty. In turn, the neural activities associated with knowledge delivery are not primarily interpreted as reward-related but as relief-related activities.

Because reward and relief are known to induce overlapping neural responses (Navratilova and Porreca, 2014; Seymour et al., 2005), our observation that the ventral striatum is activated by trivia answers in a curiosity-dependent fashion cannot arbitrate between the predictive coding and reward frameworks. Nonetheless, it must be stressed that this ventral striatal responses to knowledge delivery were not reported in previous studies (Gruber et al., 2014; Kang et al., 2009) nor observed in the second part of our trivia quizz. The most likely reason for this discrepancy is that striatal activations at the answer stage reflect relief (or reward) prediction errors: indeed, according to the temporal difference rule known to govern these activities (Pessiglione et al., 2006; Schultz et al., 1997), fully predictable reward or relief should blunt prediction error signals (O’Doherty et al., 2003; Schultz, 2013). Oppositely, when the occurrence of such outcomes is stochastic, as in the study by Jepma and colleagues (Jepma et al., 2012) and in the first run of our task, prediction error signals should be and are actually observed. Future experiments will have to manipulate answer uncertainty in a more systematic manner in order to exclude other factors that may explain the pattern of results reported here. For example, our data cannot exclude that ventral striatal responses became undetectable in the second run due to habituation, fatigue or to the involvement additional cognitive processes recruiting the ventral striatum. Besides, we observed in the second run that displaying questions which had already been answered in the first run yielded very salient activations of the striatum, possibly reflecting memory retrieval or motivational processes (Scimeca and Badre, 2012).

Yet, the scope of our study was not restricted to the relief of acute curiosity but extended to the regulation of nonspecific EC levels, which may be understood as the time-varying motivation to obtain information during the trivia quiz, independently from the specific questions displayed on screen. This lead us to formulate precise hypotheses inspired by the predictive coding framework which have no counterpart in the reward framework. In particular, the inverse relationship between EC levels and the average amount of surprise experienced during the quiz provides strong support for the idea that nonspecific EC is dynamically regulated in order to maintain uncertainty levels around an expected value. Previous research showed that curiosity, attentional allocation and associated exploratory behaviours are boosted by intermediate levels of uncertainty, and decrease whenever uncertainty becomes too high or too low (Berlyne, 1966; Gottlieb et al., 2013; Kidd and Hayden, 2015; Loewenstein, 1994). However, to our knowledge, our data is the first to demonstrate that a similar mechanism binds curiosity and uncertainty (or average surprise) across time, in a stimulus-independent fashion. Moreover, surprise mediated curiosity-driven memory encoding, hence suggesting this regulatory mechanism partly determines the maximal amount of information that individuals are able or willing to process and encode per unit of time. Furthermore, assessing the amount of surprising information at which nonspecific EC reverses from positive (i.e information attractiveness) to negative (i.e. information avoidance) may thus be useful to infer the optimal learning pace of learners.

Neuroimaging data showed that the monitoring of averaged surprise occurred primarily in the right rostrolateral PFC, in a region close but distinct from the peak coordinates previously found to control uncertainty-driven exploration (Badre et al., 2012; Boorman et al., 2009; Daw et al., 2006; Donoso et al., 2014). In this medial and anterior portion of the rlPFC, the amount of recently experienced surprise was encoded positively when participants processed new questions, whereas surprise prediction errors induced proportional deactivations of the same structure when participants were presented with new answers. Although the functional meaning of negative BOLD responses constitutes a debated topic in the neuroimaging community (Shmuel et al., 2006; Weitz et al., 2015), these deactivations are consistent with previous studies. They have been documented in the context of semantic judgments (Hayama and Rugg, 2009) and in reinforcement-learning tasks (Daw et al., 2006) where they denote exploitative rather than explorative decisions. Interestingly, at the peak coordinates previously found to control uncertainty-driven exploration (Badre et al., 2012; Boorman et al., 2009; Daw et al., 2006; Donoso et al., 2014), the encoding of surprise prediction errors was not significant at the group level, but the analysis of interindividual differences revealed a positive correlation between the negative effect of surprise on curiosity and the neural encoding of surprise prediction errors in these more posterior and lateral portions of the rlPFC. Taken together, these results suggest a possible dissociation of the neural populations within the rlPFC, from the raw monitoring of average surprise to the actual influence of this variable on EC levels.

As we have seen in the introduction, the notion of surprise is fundamental to apprehend curiosity-related behaviours and their associated neural substrates in a predictive coding perspective. This follows naturally from the fact that surprise and the information carried by any given event are actually synonymous for information theory, in which predictive coding is deeply rooted (Friston, 2010). Yet, very few EC studies systematically assessed surprise, despite its well-known empirical relationship with curiosity (Berlyne, 1962). One reason may be that surprise cannot be readily manipulated and framed as an information-theoretic or Bayesian quantity in the context of EC research as it can be done in other fields such as economic decision-making. Instead, surprise in the context of trivia quizzes and related ecological tasks is usually captured using introspective ratings, because it depends on intractable high-level representations about an open-ended world, which are themselves shaped by language. In our data, higher surprise ratings were associated with stronger vmPFC activations, likewise answers associated with more prior knowledge and new answers compared to repeated answers. This unique selectivity profile of the vmPFC makes it a strong candidate for the computation of surprise, as this signal must be derived from the comparison of new information with prior knowledge. Intriguingly, the hippocampus was not involved, despite its putative role in detecting mismatch between current observations and prior knowledge (Duncan et al., 2012; Kumaran and Maguire, 2009). However, empirical evidence supporting this hippocampal role was obtained in the presence of clear-cut expectations (e.g based on the training phase of associative learning tasks), whereas the number of possible alternatives may be too wide to be represented *a priori* in a trivia quiz. In order to disentangle the neural circuits signaling surprise as a violation of *a priori* expectations and surprise as an incongruence of new information with episodic or semantic representations retrieved *a posteriori*, it will be important to assess the presence and the accuracy of expectations when processing trivia questions in future experiment.

At this point, one may argue that our interpretation of vmPFC and rlPFC activities remains uncertain because surprise ratings correlated with curiosity, prior knowledge and interest ratings in our task. Yet, if surprise ratings indeed reflect the perceived informativeness of trivia answers, it is logical for them to covary with these variables. For example, the observed correlation between curiosity and surprise ratings — which replicates previous findings (Baranes et al., 2015) — may originate in the tight association of curiosity and selective attention (Daffner et al., 1992; Gottlieb et al., 2013) known to exacerbate surprise-related signals such as mismatch and prediction error responses (Auksztulewicz and Friston, 2015; Jiang et al., 2013). In other words, paying more attention to an answer may automatically increase the feeling of surprise it generates. The positive relationship between prior knowledge and surprise is even more straightforward, because the likelihood of ignoring an interesting fact about a movie is obviously higher when this movie has been watched rather than not. Accounting for these effects, the general linear model (GLM) used to unravel the neural correlates of surprise and surprise prediction errors included a serial orthogonalization procedure ensuring that any the variance associated with curiosity relief or prior knowledge related to answers was excluded from these regressors. This implies that the remaining variance primarily reflected unexpected information contents, as defined in the introduction. We also excluded the possibility that the surprise regressors merely reflected a valuation signal bound to each trivia item by substituting surprise for interest ratings in the GLM and by showing that the latter was not significantly associated with vmPFC activity.

One potential drawback of the current study is the number of behavioural and neural variables simultaneously considered for analysis. Although future experiments may focus on a subset of these variables in order to investigate a reduced number of processes in greater depth, we believe that the breadth of our work may help to outline a neurocognitive model of EC integrating insights from predictive coding with earlier psychological theories (Figure 6). In order to further refine and test this model, the suppressive effect of average surprise should be confirmed in other tasks probing the willingness-to-pay or the willingness-to-wait for answers (Kang et al., 2009; Marvin and Shohamy, 2016). Surprise ratings themselves should be cross-validated using implicit measures such as pupil dilation (Preuschoff et al., 2011) or eye movements (Baranes et al., 2015). Finally, it would be important to investigate the respective roles of different neuromodulatory systems in different facets of EC. For example, electrophysiological recordings in monkeys have shown that the encoding of advanced information about rewards is functionally distinct from the encoding of rewards within the vmPFC (Blanchard et al., 2015) while these processes largely overlap within the dopaminergic brainstem (Bromberg-Martin and Hikosaka, 2009; Düzel et al., 2010; Kumaran and Duzel, 2008). It would thus be tempting to propose that EC relief recruits dopaminergic neurons while surprise activates an additional neuromodulatory system. Noradrenaline would be ideally suited to mediate the interplay of prior knowledge, surprise and memory encoding highlighted by our behavioural analyses. It has indeed been involved in novelty processing (Schomaker and Meeter, 2015), long-term memory (Gibbs et al., 2010) and unexpected surprise or uncertainty signaling (Payzan-LeNestour et al., 2013; Yu and Dayan, 2005).

**Figure 6.**
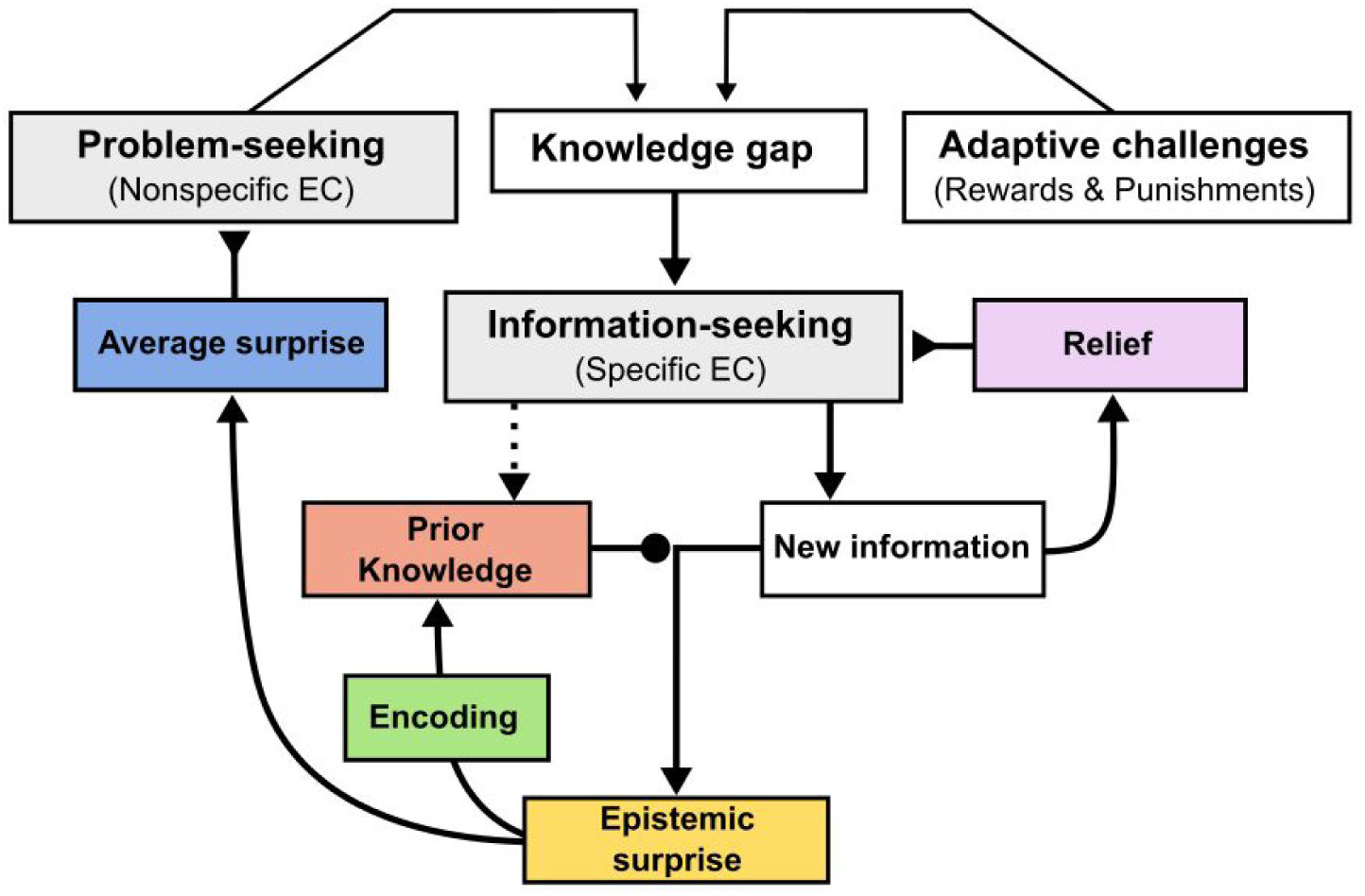
The cognitive dynamics of epistemic curiosity (EC). In this model inspired by predictive coding and earlier psychological theories (Berlyne, 1966; Loewenstein, 1994), all information-seeking behaviours start with the awareness of a “knowledge gap” encountered while solving adaptive challenges or seeking new epistemic problems. Here, specific EC corresponds to internal (i.e recollecting one’s own knowledge) or external (i.e exploring the environment) information-seeking behaviours engaged to gather new information and reduce the uncertainty bound to knowledge gaps. The comparison of new information with prior knowledge may then elicit epistemic surprise, thereby facilitating memory encoding. The key hypothesis of this model is that the human cognitive system represents the average amount of surprise experienced in a given context and that this variable exerts a suppressive effect on the nonspecific component of EC which underlies problem-seeking behaviours. Problem-seeking behaviours — a hallmark of most intellectual, artistic and scientific professions — may thus derive from a disruption of this process. Conversely, the lack of curiosity may derive from its exacerbation.

To conclude, understanding the regulation of EC and its neural implementation in the human brain will require intense research efforts in domains as diverse as memory, attention, linguistics and decision-making. This endeavour must build on the complementary insights provided by the reinforcement-learning and predictive coding frameworks, as well as other information-seeking principles such as learning progress maximization (Gottlieb et al., 2013). Taking into account the dynamical relationship between surprise and curiosity may help integrate this diverse literature and open the path to new strategies of knowledge transfer in the classroom.

## Materials and Methods

See SI Materials and Methods for complete details regarding participants, stimuli, tasks, preprocessing and data analysis.

### Participants

Twenty-two right-handed students (11 females, 11 males; mean age: 22.9; range: 19-28) were recruited through advertisements in an art cinema and via university mailing lists. This sample size matched the range of existing neuroimaging studies on epistemic curiosity (Gruber et al., 2014; Kang et al., 2009). No participant was excluded from data analyses. All were paid at the fixed rate of 60€ for their participation in the study. A few days before the experimental session, participants signed informed consent after exhaustive explanations were provided. The entire protocol was approved by the local ethics committee of Sud-Est II, France (authorization number: 2011-056-2).

### Stimuli

Sixty question-answer pairs were selected amongst the 120 pre-screened trivia items (Fig. S3 and Table S1). For the prior knowledge localizer task, we used personalized sets of movie titles, different from those encountered in the main trivia task.

### Time course of the experiment

At their arrival to the MRI lab, participants were reminded that they would be exposed to cinema-related trivia questions and warned that those questions had been selected for being interesting but rarely known, even to cinema lovers. In the first functional run, each trial started with a jittered fixation cross. Then, participants had to read one of the 60 pre-screened trivia questions and to signal end of reading with a button press. After a fixed interval, a continuous gauge appeared. Participants had then to rate their curiosity by keeping the left button pressed until the gauge reach the desired point. Another jittered fixation cross preceded the delivery of either an answer (50% of the trials) or hashtags. The temporal order of items was randomized for each participant independently. In the second functional run, participants were verbally instructed that they would be presented again with all the questions, and that this time they would simply have to indicate whether the correct answer came spontaneously to their mind or not. All questions were again preceded and followed by a fixation cross. In this second run, answers were delivered in all trials. The temporal order of items was re-randomized for each participant independently. In the third functional run, participants were presented with 30 watched movie titles, 30 unwatched movie titles, and 30 hashtags “#”. When a target was appeared on the screen participants had simply to report whether they had seen this movie title or not. Once outside the scanner, participants were first presented with an unexpected memory test in which they had to write down the answer of the 60 trivia questions encountered in the task. They also reported which answers they were expecting (13.2±8.8%) or knew already for sure (4.4±5.1%) before the task. Then, all questions and answers were shown together, and participants were asked to rate their surprise levels (from 1 “not at all” to 5 “yes, a lot”) and to report the thirty items they found the most interesting. To conclude, they filled an epistemic curiosity questionnaire (Litman and Spielberger, 2003) designed to capture specific (i.e deprivation) and diversive (i.e interest) EC.

### Behavioural analyses

The modelling of nonspecific EC levels as a function of epistemic surprise used a delta-rule of the form *Q*_*t*+1_ = *Q*_*t*_ + α(*R* − *Q*_*t*_), where Q is initialized at 0 and updated on each trial by the prediction error term R-Q, times a learning rate α. In the best-fitted model termed Q{sur},the delivery of an answer was coded as R=S while the absence of answer was coded as R=0, with S corresponding to the surprise rating given by the participant for that particular item. In order to ascertain that this approach was useful to explain variance in curiosity ratings, we compared a range of alternative models using a Bayesian group comparison approach (Fig. 4A, Fig. S1A-D). All details regarding model fit, model comparison and mediation analyses can be found in SI Materials and Methods. When testing correlation or difference between two variables, parametric statistical tests (i.e Pearson correlation coefficient and Student t-test) were used when the two variables were normally distributed. Their non-parametric equivalent was used otherwise (i.e Spearman rank correlation and Wilcoxon u-test).

### fMRI analyses

Statistical analyses of fMRI signals were performed using a conventional two-levels random-effects approach with SPM8. All general linear models (GLM) included 6 unconvolved motion parameters obtained from the realignment step, in order to covary out potential movement-related artifacts in the BOLD signal. All regressors of interest were convolved with the canonical hemodynamic response function (HRF). All GLM models included a high-pass filter to remove low-frequency artifacts from the data as well as a run-specific intercept. All motor responses recorded were modeled using a zero-duration Dirac function. Voxel-wise thresholds used to generate SPM maps were either p<0.005^UNC^ (parametric contrasts) or p<0.001^UNC^ (categorical contrasts), unless notified otherwise. All statistical inferences based on whole-brain analyses met the standard multiple comparison threshold (p<0.05^FWE^) at the cluster level.

In the first run (GLM1), the question, rating and outcome stages were modeled separately using boxcar functions set to the duration of each individual event. Questions for which the participant did not know the answer were parametrically modulated by four regressors, orthogonalized in the following order: Qsur (average surprise), Prior knowledge, Curiosity, Subsequent recall. At the outcome stage, answers and hashtags were also parametrically modulated using four regressors, orthogonalized in the following order: Curiosity, Prior knowledge, Surprise prediction error (or Surprise), Subsequent recall. Questions and answers for which participants knew the answer before starting the experiment were modeled separately and not included in any contrast. In the second run, we splitted questions and answers regressors as a function of their status in the first run (i.e. items answered or not in run 1) and participants’ ability to recall spontaneously the answer or not. This resulted in two “HIT” regressors (items previously answered and remembered, at the question and answer stages) and two “correct rejection” (CR) regressors (unanswered and correctly classified as such, also at both stages). Questions (HIT and CR) and answers were parametrically modulated using 4 regressors, orthogonalized in the following order: Curiosity, Prior Knowledge, Surprise, Subsequent recall. In the third run, we modelled the onset of hashtags, watched movies and unwatched movies separately using zero-duration Dirac functions. Note that Table S2 describe all the variables used a regressors in both behavioural and fMRI analyses.

Concerning ROI analyses, the mask used to extract effects from the peaks previously reported in the literature study the contribution of the rlPFC to uncertainty-driven exploration were 3mm-radius spheres centered around the MNI coordinates reported in the original papers (explicitly displayed on Fig. 4b). For the multiple ROIs analyses reported Fig. 5 and Fig. S1b-h, we used the following approach: (i) clusters surviving a voxel-wise threshold of p<0.05FWE were extracted from the [new answer>hashtag] contrast (run 1; dlPFC, vmPFC, HPC, STS, Precuneus, IPL), (ii) clusters surviving a cluster-wise threshold of p<0.05FWE (voxel-wise threshold: p<0.005unc) were extracted from the parametric curiosity contrasts at the question (dmPFC, IPL) and answer (ventral striatum) stages of run 1. For each of the 8 regions, the mirror (x-flipped) ROI was added to the mask itself, so that every ROIs were strictly symmetric and identical across the two hemispheres. The peak MNI coordinates corresponding to previous studies investigating the uncertainty-dependent exploration were: [27 57 6] for Daw et al. (Daw et al., 2006), [36, 54, 0] for Boorman et al. (Boorman et al., 2009), [32 56 12] for Donoso et al. (Donoso et al., 2014), 4: [35 56 −8] for Badre et al. (Badre et al., 2012). Peristimulus time-course histograms (PSTH, sampled at 1Hz) were computed using the toolbox *rfxplot* for Matlab (Gläscher, 2009). These time-decomposed effects were thus re-estimated using the first eigenvariate extracted from the regions of interest, after adjustment for run intercept and movement-related variance.

## ACKNOWLEDGEMENTS

We are grateful to the staff of CERMEP–Imagerie du Vivant for helpful assistance with data collection and to the movie theater Comoedia (Lyon, France) for offering us the opportunity to perform the behavioural experiment in their premises. We must also thank Giulia Gennari and Roshan Cools for their precious comments on earlier versions of the manuscript. Finally, we would like to thank all participants for their curiosity in our study.

## AUTHOR CONTRIBUTIONS

R.L and T.M designed and performed the experiments. R.L and T.M analyzed the behavioural data. R.L analyzed the fMRI data. R.L and T.M wrote the paper. M.M funded the experiments and participated in writing the paper.

## References

Ahsan, R.L., Allom, R., Gousias, I.S., Habib, H., Turkheimer, F.E., Free, S., Lemieux, L., Myers, R., Duncan, J.S., Brooks, D.J., Koepp, M.J., Hammers, A., 2007. Volumes, spatial extents and a probabilistic atlas of the human basal ganglia and thalamus. Neuroimage 38, 261–270.

Auksztulewicz, R., Friston, K., 2015. Attentional Enhancement of Auditory Mismatch Responses: a DCM/MEG Study. Cereb. Cortex 25, 4273–4283.

Badre, D., Doll, B.B., Long, N.M., Frank, M.J., 2012. Rostrolateral prefrontal cortex and individual differences in uncertainty-driven exploration. Neuron 73, 595–607.

Baranes, A., Oudeyer, P.-Y., Gottlieb, J., 2015. Eye movements reveal epistemic curiosity in human observers. Vision Res. 117, 81–90.

Bateman, T.S., Hess, A.M., 2015. Different personal propensities among scientists relate to deeper vs. broader knowledge contributions. Proc. Natl. Acad. Sci. U. S. A. 112, 3653–3658.

Berlyne, D.E., 1966. Curiosity and exploration. Science 153, 25–33.

Berlyne, D.E., 1962. Uncertainty and epistemic curiosity. Br. J. Psychol. 53, 27–34.

Berlyne, D.E., 1954. A theory of human curiosity. Br. J. Psychol. 45, 180–191.

Blanchard, T.C., Hayden, B.Y., Bromberg-Martin, E.S., 2015. Orbitofrontal cortex uses distinct codes for different choice attributes in decisions motivated by curiosity. Neuron 85, 602–614.

Boorman, E.D., Behrens, T.E.J., Woolrich, M.W., Rushworth, M.F.S., 2009. How green is the grass on the other side? Frontopolar cortex and the evidence in favor of alternative courses of action. Neuron 62, 733–743.

Bromberg-Martin, E.S., Hikosaka, O., 2009. Midbrain dopamine neurons signal preference for advance information about upcoming rewards. Neuron 63, 119–126.

Clark, A., 2013. Whatever next? Predictive brains, situated agents, and the future of cognitive science. Behav. Brain Sci. 36, 181–204.

Daffner, K.R., Scinto, L.F., Weintraub, S., Guinessey, J.E., Mesulam, M.M., 1992. Diminished curiosity in patients with probable Alzheimer’s disease as measured by exploratory eye movements. Neurology 42, 320–328.

Daw, N.D., O’Doherty, J.P., Dayan, P., Seymour, B., Dolan, R.J., 2006. Cortical substrates for exploratory decisions in humans. Nature 441, 876–879.

Donoso, M., Collins, A.G.E., Koechlin, E., 2014. Human cognition. Foundations of human reasoning in the prefrontal cortex. Science 344, 1481–1486.

Duncan, K., Ketz, N., Inati, S.J., Davachi, L., 2012. Evidence for area CA1 as a match/mismatch detector: a high-resolution fMRI study of the human hippocampus. Hippocampus 22, 389–398.

Düzel, E., Bunzeck, N., Guitart-Masip, M., Düzel, S., 2010. NOvelty-related motivation of anticipation and exploration by dopamine (NOMAD): implications for healthy aging. Neurosci. Biobehav. Rev. 34, 660–669.

Friston, K., 2010. The free-energy principle: a unified brain theory? Nat. Rev. Neurosci. 11, 127–138.

Friston, K., Rigoli, F., Ognibene, D., Mathys, C., Fitzgerald, T., Pezzulo, G., 2015. Active inference and epistemic value. Cogn. Neurosci. 6, 187–214.

Friston, K., Thornton, C., Clark, A., 2012. Free-energy minimization and the dark-room problem. Front. Psychol. 3, 130.

Garrido, M.I., Barnes, G.R., Kumaran, D., Maguire, E.A., Dolan, R.J., 2015. Ventromedial prefrontal cortex drives hippocampal theta oscillations induced by mismatch computations. Neuroimage 120, 362–370.

Garrison, J., Erdeniz, B., Done, J., 2013. Prediction error in reinforcement learning: a meta-analysis of neuroimaging studies. Neurosci. Biobehav. Rev. 37, 1297–1310.

Ghosh, V.E., Moscovitch, M., Melo Colella, B., Gilboa, A., 2014. Schema representation in patients with ventromedial PFC lesions. J. Neurosci. 34, 12057–12070.

Gibbs, M.E., Hutchinson, D.S., Summers, R.J., 2010. Noradrenaline release in the locus coeruleus modulates memory formation and consolidation; roles for α- and β-adrenergic receptors. Neuroscience 170, 1209–1222.

Gigerenzer, G., Garcia-Retamero, R., 2017. Cassandra’s regret: The psychology of not wanting to know. Psychol. Rev. 124, 179–196.

Gläscher, J., 2009. Visualization of group inference data in functional neuroimaging. Neuroinformatics 7, 73–82.

Gottlieb, J., Oudeyer, P.-Y., Lopes, M., Baranes, A., 2013. Information-seeking, curiosity, and attention: computational and neural mechanisms. Trends Cogn. Sci. 17, 585–593.

Gruber, M.J., Gelman, B.D., Ranganath, C., 2014. States of curiosity modulate hippocampus-dependent learning via the dopaminergic circuit. Neuron 84, 486–496.

Hayama, H.R., Rugg, M.D., 2009. Right dorsolateral prefrontal cortex is engaged during post-retrieval processing of both episodic and semantic information. Neuropsychologia 47, 2409–2416.

Hebb, 1955. Drives and the C.N.S. (conceptual nervous system). Psychol. Rev. 62, 243.

Hirsh, J.B., Mar, R.A., Peterson, J.B., 2012. Psychological entropy: a framework for understanding uncertainty-related anxiety. Psychol. Rev. 119, 304–320.

Hsee, C.K., Ruan, B., 2016. The Pandora Effect: The Power and Peril of Curiosity. Psychol. Sci. 27, 659–666.

Jepma, M., Verdonschot, R.G., van Steenbergen, H., Rombouts, S.A.R.B., Nieuwenhuis, S., 2012. Neural mechanisms underlying the induction and relief of perceptual curiosity. Front. Behav. Neurosci. 6, 5.

Jiang, J., Summerfield, C., Egner, T., 2013. Attention sharpens the distinction between expected and unexpected percepts in the visual brain. J. Neurosci. 33, 18438–18447.

Kang, M.J., Hsu, M., Krajbich, I.M., Loewenstein, G., McClure, S.M., Wang, J.T.-Y., Camerer, C.F., 2009. The wick in the candle of learning: epistemic curiosity activates reward circuitry and enhances memory. Psychol. Sci. 20, 963–973.

Kashdan, T.B., Gallagher, M.W., Silvia, P.J., Winterstein, B.P., Breen, W.E., Terhar, D., Steger, M.F., 2009. The Curiosity and Exploration Inventory-II: Development, Factor Structure, and Psychometrics. J. Res. Pers. 43, 987–998.

Kidd, C., Hayden, B.Y., 2015. The Psychology and Neuroscience of Curiosity. Neuron 88, 449–460.

Kumaran, D., Duzel, E., 2008. The hippocampus and dopaminergic midbrain: old couple, new insights. Neuron 60, 197–200.

Kumaran, D., Maguire, E.A., 2009. Novelty signals: a window into hippocampal information processing. Trends Cogn. Sci. 13, 47–54.

Lebreton, M., Abitbol, R., Daunizeau, J., Pessiglione, M., 2015. Automatic integration of confidence in the brain valuation signal. Nat. Neurosci. 18, 1159–1167.

Lighthall, N.R., Huettel, S.A., Cabeza, R., 2014. Functional compensation in the ventromedial prefrontal cortex improves memory-dependent decisions in older adults. J. Neurosci. 34, 15648–15657.

Litman, J.A., 2010. Relationships between measures of I- and D-type curiosity, ambiguity tolerance, and need for closure: An initial test of the wanting-liking model of information-seeking. Pers. Individ. Dif. 48, 397–402.

Litman, J.A., Spielberger, C.D., 2003. Measuring epistemic curiosity and its diversive and specific components. J. Pers. Assess. 80, 75–86.

Litman, J., Hutchins, T., Russon, R., 2005. Epistemic curiosity, feeling-of-knowing, and exploratory behaviour. Cognition and Emotion 19, 559–582.

Loewenstein, G., 1994. The psychology of curiosity: A review and reinterpretation. Psychol. Bull. 116, 75.

Marvin, C.B., Shohamy, D., 2016. Curiosity and reward: Valence predicts choice and information prediction errors enhance learning. J. Exp. Psychol. Gen. 145, 266–272.

McGillivray, S., Murayama, K., Castel, A.D., 2015. Thirst for knowledge: The effects of curiosity and interest on memory in younger and older adults. Psychol. Aging 30, 835–841.

Navratilova, E., Porreca, F., 2014. Reward and motivation in pain and pain relief. Nat. Neurosci. 17, 1304–1312.

O’Doherty, J.P., Dayan, P., Friston, K., Critchley, H., Dolan, R.J., 2003. Temporal difference models and reward-related learning in the human brain. Neuron 38, 329–337.

Payzan-LeNestour, E., Dunne, S., Bossaerts, P., O’Doherty, J.P., 2013. The neural representation of unexpected uncertainty during value-based decision making. Neuron 79, 191–201.

Pessiglione, M., Seymour, B., Flandin, G., Dolan, R.J., Frith, C.D., 2006. Dopamine-dependent prediction errors underpin reward-seeking behaviour in humans. Nature 442, 1042–1045.

Pierce, J.P., Distefan, J.M., Kaplan, R.M., Gilpin, E.A., 2005. The role of curiosity in smoking initiation. Addict. Behav. 30, 685–696.

Preuschoff, K., ’t Hart, B.M., Einhäuser, W., 2011. Pupil Dilation Signals Surprise: Evidence for Noradrenaline’s Role in Decision Making. Front. Neurosci. 5, 115.

Raja Beharelle, A., Polanía, R., Hare, T.A., Ruff, C.C., 2015. Transcranial Stimulation over Frontopolar Cortex Elucidates the Choice Attributes and Neural Mechanisms Used to Resolve Exploration–Exploitation Trade-Offs. J. Neurosci. 35, 14544–14556.

Rissman, J., Chow, T.E., Reggente, N., Wagner, A.D., 2016. Decoding fMRI Signatures of Real-world Autobiographical Memory Retrieval. J. Cogn. Neurosci. 28, 604–620.

Schomaker, J., Meeter, M., 2015. Short- and long-lasting consequences of novelty, deviance and surprise on brain and cognition. Neurosci. Biobehav. Rev. 55, 268–279.

Schultz, W., 2013. Updating dopamine reward signals. Curr. Opin. Neurobiol. 23, 229–238.

Schultz, W., Dayan, P., Montague, P.R., 1997. A neural substrate of prediction and reward. Science 275, 1593–1599.

Schwartenbeck, P., FitzGerald, T., Dolan, R., Friston, K., 2013. Exploration, novelty, surprise, and free energy minimization. Front. Psychol. 4, 710.

Scimeca, J.M., Badre, D., 2012. Striatal contributions to declarative memory retrieval. Neuron 75, 380–392.

Seymour, B., O’Doherty, J.P., Koltzenburg, M., Wiech, K., Frackowiak, R., Friston, K., Dolan, R., 2005. Opponent appetitive-aversive neural processes underlie predictive learning of pain relief. Nat. Neurosci. 8, 1234–1240.

Shmuel, A., Augath, M., Oeltermann, A., Logothetis, N.K., 2006. Negative functional MRI response correlates with decreases in neuronal activity in monkey visual area V1. Nat. Neurosci. 9, 569–577.

Sweeny, K., Melnyk, D., Miller, W., Shepperd, J.A., 2010. Information avoidance: Who, what, when, and why. Rev. Gen. Psychol. 14, 340.

van Dijk, E., Zeelenberg, M., 2007. When curiosity killed regret: Avoiding or seeking the unknown in decision-making under uncertainty. J. Exp. Soc. Psychol. 43, 656–662.

van Kesteren, M.T.R., Beul, S.F., Takashima, A., Henson, R.N., Ruiter, D.J., Fernández, G., 2013. Differential roles for medial prefrontal and medial temporal cortices in schema-dependent encoding: from congruent to incongruent. Neuropsychologia 51, 2352–2359.

Von Stumm, S., Hell, B., Chamorro-Premuzic, T., 2011. The hungry mind intellectual curiosity is the third pillar of academic performance. Perspect. Psychol. Sci. 6, 574–588.

Weitz, A.J., Fang, Z., Lee, H.J., Fisher, R.S., Smith, W.C., Choy, M., Liu, J., Lin, P., Rosenberg, M., Lee, J.H., 2015. Optogenetic fMRI reveals distinct, frequency-dependent networks recruited by dorsal and intermediate hippocampus stimulations. Neuroimage 107, 229–241.

Yu, A.J., Dayan, P., 2005. Uncertainty, neuromodulation, and attention. Neuron 46, 681–692.

