## Supplementary Materials for "The control of epistemic curiosity in the human brain"

**SUPPLEMENTARY INFORMATION**

**Participants**

Twenty-two right-handed students (11 females, 11 males; mean age: 22.9; range: 19-28) were recruited through advertisements in an art cinema and via university mailing lists. This sample size matched the range of existing neuroimaging studies on epistemic curiosity (Gruber et al., 2014; Kang et al., 2009). No participant was excluded from data analyses. All were paid at the fixed rate of 60€ for their participation in the study. A few days before the experimental session, participants signed informed consent after exhaustive explanations were provided. They were also given a list of 215 movie titles and asked to indicate for each of them to what extent they knew the movie (from 1= *never heard of it* to 4 = *seen it several times*). Target movie titles were covertly included in this list, which enabled us to quantify prior knowledge about trivia items (i.e. watched/unwatched status). For the behavioural experiment performed to select and validate the trivia used in the fMRI study, 64 participants of all ages were invited to complete a computerized evaluation of candidate trivia items (Supp. Fig. S3A) after filing a consent form. The behavioural experiment took place in an art cinema (Comoedia, Lyon). After completing the task (about 20 minutes), they were offered to pick a book among a large selection of novels and essays. The entire protocol was approved by the local ethics committee of Sud-Est II, France (authorization number: 2011-056-2).

**Stimuli**

Sixty question-answer pairs were selected amongst the 120 pre-screened trivia items (Supp. Fig. S3A and Table S1). The trivia questions included in the fMRI experiment were chosen to maximize reported surprise, interest and knowledge about the target movies (Supp. Fig S3B). In addition, items were selected and designed to minimize the chances that participants would know or guess the answers. In the fMRI experiment, the frequency of known answers was therefore very low (5.5±6.1%: range: 0-18%) and known items were always modeled separately and excluded from all analyses (except for Supp. Fig. S1C). Moreover, we ensured that items associated with the two main conditions (i.e. items answered or not in the first run) were highly matched for characters count (for both questions and answers), curiosity, surprise and interest (all $p>0.75$). Finally, we counterbalanced across participants the subsets of items associated with each condition.

For the prior knowledge localizer task, we used the 215-items questionnaire to create two personalized sets of movie titles, different from those encountered in the main trivia task: 30 watched movies (if possible, watched less than two years before the experimental session) and 30 unwatched movies (if possible, with titles known). For two participants who had not seen enough movies in the list, we included movies seen more than two years before within the pool of watched movies (10 and 26 items, respectively).

**Time course of the fMRI experiment**

At their arrival to the MRI lab, participants were reminded that they would be exposed to cinema-related trivia questions and warned that those questions had been selected for being interesting but rarely known, even to cinema lovers. Once in the scanner, they completed sixteen training items to improve self-calibration in curiosity ratings (those items were not redundant with those of the main task) after receiving the following instructions (hereafter translated from French):

*You are about to begin an experiment about intellectual curiosity and cinema in the specific conditions of the MRI scanner.*

*[Slide 1] During the calibration of the scanner and the acquisition of the anatomical image of our brain, you will practice the task that you will be doing while we will record your cerebral activities. This training must in particular enable you to manipulate properly the response gauge with which you will indicate to what extent you are curious to know the answers to the questions we are going to present you.*

*[Slide 2, showing a fixation cross] Each trial will begin with a small symbol signaling that a question is about to appear.*

*[Slide 3, showing a dummy trivia question] After a few seconds, the question will appear. Take the time to read it properly. Once you have read it, press the left button (index).*

*[Slide 4, showing the gauge and the question] Once you press the button, the curiosity gauge will appear. By keeping the left button pressed, you can increase the gauge up to how much you are curious.*

*[Slide 5, showing the gauge and the question] If you are sure to know the answer, press the right button (major finger) when the answer comes to your mind. NB: if you think you know the answer but don't remember it (answer on the tip of the tongue), don't answer with the right finger but indicate your curiosity level.*

*[Slide 6] Once you have raised the gauge up to the level corresponding to your curiosity, a fixation cross will appear on the screen.*

*[Slide 7, showing the answer to the dummy question] Finally, the answer to the question will be displayed during a few seconds. However, during the first part of the experiment, we will only delivered 50% of the answers. NB: there is NO relationship between your curiosity rating and the likelihood of of receiving or not the answer.*

In the first functional run, each trial started with a jittered fixation cross (exponential distribution; mean: 4.2s: range: 3-7.5 seconds). Then, participants had to read one of the 60 pre-screened trivia questions and to signal end of reading with a button press (right index finger; average reading time: 5.1±1.49s). After a fixed interval of 750ms, a continuous gauge appeared. Participants had then to use their index finger to rate their curiosity by keeping the left button pressed until the gauge reach the desired point (maximum curiosity: 2.5s). In case they would know the answer already, they were instructed to answer with the right finger and then had to wait for 2s. Another jittered fixation cross (exponential distribution; mean: 4.2s: range: 3-7.5s) preceded the delivery of either an answer (50% of the trials) or hash tags "#" (3s, fixed duration). The temporal order of items was randomized for each participant independently. Importantly, we checked that the multiplication of parametric regressors did not induce major change in the statistics reported in Fig. 3 to 5. All effects remained significant when the different parametric regressors were tested in separate GLM, except for the curiosity-dependent modulation of ventral striatal activity reported in Fig. 3A. However, given its previous involvement in epistemic curiosity, we decided to maintain this results in the manuscript as it was significant in an anatomical mask of the nucleus accumbens (Fig. 3B) and survived small-volume correction using an anatomical mask of the whole striatum (data not shown, pFWE<0.05, SVC).

In the second functional run, participants were verbally instructed that they would be presented again with all the questions, and that this time they would simply have to indicate whether the correct answer came spontaneously to their mind or not (average response time: 3.7±0.83s). To do so, they had to select either a "light bulb" or a "cloud" associated with each situation, respectively (black and white drawings of similar size displayed on the left and right of the question; side counterbalanced across trials). All questions were again preceded and followed by a fixation cross (exponential distribution; mean: 4.2s: range: 3-7.5s). In this second run, answers were delivered in all trials (3s, fixed duration). The temporal order of items was re-randomized for each participant independently. Importantly, we checked that the multiplication of parametric regressors did not induce major changes in the statistics reported in Fig. 3 to 5. All effects remained significant when the different parametric regressors were tested in separate GLM.

In the third functional run, participants were presented with 30 watched movie titles, 30 unwatched movie titles, and 30 hashtags "#". Each trial began with a fixation cross (mean: 2.5s: range: 2-6s). Then a target was appeared on the screen for a fixed duration (3s) together with two dots, associated with the mentions "seen" and "unseen" (on the left and right of the movie title) or "skip" (on both side, in case of hash tags). The side of "seen" and "unseen" mentions was counterbalanced across trials and the temporal order of items was randomized for each participant independently.

Once outside the scanner, participants were first presented with an unexpected memory test in which they had to write down the answer of the 60 trivia questions encountered in the task. At this stage, they also reported which answers they were expecting (13.2±8.8%) or knew already for sure (4.4±5.1%) before the task. Then, all questions and answers were shown together, and participants were asked to rate their surprise levels (from 1 "not at all" to 5 "yes, a lot") and to report the thirty items they found the most interesting. To conclude, they filled an epistemic curiosity questionnaire (Litman and Spielberger, 2003) designed to capture specific (i.e deprivation) and diversive (i.e interest) EC. All behavioural tasks were programmed using Presentation (www.neurobs.com).

**fMRI acquisition**

Imaging was conducted on a Siemens Sonata scanner (1.5T), using an eight-channel head coil. Twenty six interleaved slices tilted relative to the anterior commissure – posterior commissure line (20-30°) were acquired per volume. We acquired an average of 837 echo-planar T2*-weighted functional volumes per subject (TR = 2.5; TE = 60 ms; FOV = 220 mm; matrix = 64 x 64; voxel size = 3.4 x 3.4 x 4mm). Following the fMRI session, a high-resolution T1-weighted anatomical scan was acquired. Before the functional acquisition, a gradient-field map was acquired using a gradient echo sequence and was applied for distortion-correction of the acquired functional images in order to improve local field homogeneity and minimize susceptibility artifacts, for example in the ventral parts of the prefrontal cortex.

**fMRI preprocessing**

All preprocessing steps were performed using SPM8. The first four volumes of each run were removed to allow for T1 equilibrium effects. For each participant, functional images were time-corrected, realigned, unwarped using the magnitude and phase images, and coregistered to the anatomical scan. The six movement parameters were derived from the iterative realignment procedure carried out by SPM8 (three for translation, three for rotation). The anatomical scan was then normalized to the MNI space using the ICBM152 template brain and the resulting non-linear transformation matrix was applied to the functional images. Finally, the normalized functional images were spatially smoothed with an 8 mm Gaussian kernel.

**fMRI analyses**

Statistical analyses of fMRI signals were performed using a conventional two-levels random-effects approach with SPM8. All general linear models (GLM) described below included the 6 unconvolved motion parameters from the realignment step, in order to covary out potential movement-related artifacts in the BOLD signal. All regressors of interest were convolved with the canonical hemodynamic response function (HRF). All GLM models included a high-pass filter to remove low-frequency artifacts from the data (cut-off = 128s) as well as a run-specific intercept. Temporal autocorrelation was modeled using an AR(1) process. All motor responses recorded were modeled using a zero-duration Dirac function. Voxel-wise thresholds used to generate SPM maps were either $p<0.005^{UNC}$ (parametric contrasts) or $p<0.001^{UNC}$ (categorical contrasts), unless notified otherwise. All statistical inferences based on whole-brain analyses satisfied the standard multiple comparison threshold ($p<0.05^{FWE}$) at the cluster level.

*GLM models*

In the first run (GLM1), the question, rating and outcome stages were modeled separately using boxcar functions set to the duration of each individual event. This decision to use boxcars was justified by an analysis of the residuals produced by the GLMs at the first level, compared with those from the homologous model using Dirac functions (difference in log-likelihood (LL) against homologous Dirac model: 271.7). Questions for which the participant did not know the answer were parametrically modulated by four regressors, orthogonalized in the following order:

1°) Qsur: value of the surprise accumulator (see "behavioural analyses" section, below).

2°) Prior knowledge: 1 if target movie title had been watched by the participant, 0 otherwise.

3°) Curiosity: value from 0 (excluded) to 1 (maximum curiosity).

4°) Subsequent recall: 1 if item subsequently recalled, 0 otherwise.

At the outcome stage, answers and hashtags were also parametrically modulated using four regressors, orthogonalized in the following order:

1°) Curiosity.

2°) Prior knowledge.

3°) Surprise prediction error or Surprise (see below).

4°) Subsequent recall

Questions and answers for which participants knew the answer before starting the experiment were modeled separately and not included in any contrast, except for the contrast reported Fig. S1C. In order to uncover the neural correlates of surprise, surprise ratings were simply substituted to surprise prediction errors, keeping all other aspects of the analysis identical.

In the second run, questions and answers were both modeled using Dirac functions.. Again, this decision was principled by the analysis of first-level residuals (difference in LL against homologous boxcar model: 23.0). We splitted questions and answers regressors as a function of their status in the first run (i.e. items answered or not in run 1) and participants' ability to recall spontaneously the answer or not. This resulted in two "HIT" regressors (items previously answered and remembered, at the question and answer stages) and two "correct rejection" (CR) regressors (unanswered and correctly classified as such, also at both stages). Questions (HIT and CR) were parametrically modulated using 4 regressors, orthogonalized in the following order:

1°) Curiosity

2°) Prior Knowledge

3°) Surprise

4°) Subsequent recall

Answers (HIT and CR) were also modulated using 4 regressors, orthogonalized in the following order:

1°) Curiosity

2°) Prior Knowledge

3°) Surprise

4°) Subsequent recall

Items which had been answered in the first run but could not be spontaneously recalled by the participants were modeled separately (MISS regressors). Items which were already known before starting the experiment were also modeled separately and not included in any analysis.

In the third run, we modelled the onset of hashtags, watched movies and unwatched movies separately using zero-duration Dirac functions. Given the short duration of each trial, we lowered the cut-off of the high-pass filter (64s instead of 128s).

*ROI and PSTH analyses*

Concerning ROI analyses, the mask used to extract effects from the peaks previously reported in the literature study the contribution of the rlPFC to uncertainty-driven exploration were 3mm-radius spheres centered around the MNI coordinates reported in the original papers (explicitly displayed on Fig. 4b). For the multiple ROIs analyses reported Fig 3C, Fig. 5B-F and Supp. Fig. S1B-H, we used the following method: (i) clusters surviving a voxel-wise threshold of p<0.05FWE were extracted from the [new answer>hashtag] contrast (run 1; dlPFC, vmPFC, HPC, STS, Precuneus), (ii) clusters surviving a cluster-wise threshold of p<0.05FWE (voxel-wise threshold: p<0.005unc) were extracted from the parametric curiosity contrasts at the question (dmPFC, IPL) and answer (ventral striatum) stages of run 1. For each of the 8 regions, the mirror (x-flipped) ROI was added to the mask itself, so that every ROIs were strictly symmetric and identical across the two hemispheres. Finally, the nucleus accumbens mask (Fig. 3B) was obtained from an anatomical probabilistic atlas of the basal ganglia (Ahsan et al., 2007).

Peristimulus time-course histograms (PSTH, sampled at 1Hz) were computed using the toolbox *rfxplot* for Matlab (Gläscher, 2009). These time-decomposed effects were thus re-estimated using the first eigenvariate extracted from the regions of interest, after adjustment for run intercept and movement-related variance.

When testing correlation or difference between two variables, parametric statistical tests (i.e Pearson correlation coefficient and Student t-test) were used when the two variables were normally distributed. Their non-parametric equivalent was used otherwise (i.e Spearman rank correlation and Wilcoxon u-test).

**behavioural analyses**

*Delta-rule*

The modelling of nonspecific EC levels as a function of epistemic surprise used the following delta-rule:

$$Q_{t+1} = Q_t + \alpha(R - Q_t) \quad \text{(Equation 1)}$$

where Q is initialized at 0 and updated on each trial by the prediction error term R-Q, times a learning rate α. In the most simple model termed Q{0-1}, the delivery of an answer was coded as R=1 while the absence of answer was coded as R=0, so that the variable Q represents the amount of knowledge recently delivered to the participant, which enabled us to explore whether knowledge tended to reinforce or saturate curiosity over time. In the best-fitted model termed Q{sur},the delivery of an answer was coded as R=S while the absence of answer was coded as R=0, with S corresponding to the surprise rating given by the participant for that particular item.

In order to ascertain that this approach was useful to explain variance in curiosity ratings, we compared a range of alternative models using a Bayesian group comparison approach (Fig. 4A, Fig. S2A-D), as implemented in the toolbox VBA (Daunizeau et al., 2014) for Matlab (http://mbb-team.github.io/VBA-toolbox/). Alternative models were: Q{0-1}, Q{sur}, time, Q{0-1} & time, Q{sur} & time. The "time" model was used to ascertain that our delta-rule was not merely capturing a linear (increasing or decreasing) trend in curiosity ratings but rather an information-dependent process. An intercept was included in all models. Learning rates were treated as a fixed-effect in order to limit model complexity and facilitate the interpretation of individual differences and correlates of surprise accumulation. Subject-level estimations were performed using the *fitglm* algorithm provided in Matlab. When the fit was performed on continuous curiosity ratings, we assumed that those were normally distributed. However, because this assumption was violated in 4 participants, we also check that the same results could be observed using binarized curiosity levels (ie. superior or inferior to 50%, corresponding to the half-maximum of the curiosity gauge).

*Generalized Estimating Equations*

To confirm the complementary contributions of prior knowledge, surprise and curiosity in facilitating recall performance, we performed a Generalized Estimating Equations analysis (GEE) analysis, as implemented by SPSS 21 (https://www.ibm.com/analytics/us/en/technology/spss/). Successful recall was coded as 1 and unsuccessful recall as 0 and predicted by mean of a logistic regression. The analysis included a participant-specific intercept, trivia id as a within-participant effects, and curiosity, prior knowledge and surprise as random effects.

*Mediation analysis*

The multi-level mediation analysis (Fig. 2D) was performed using the Mediation toolbox[26] for Matlab (https://github.com/canlab/MediationToolbox). Curiosity and surprise levels were z-scored for each participant separately, after removing items which were known to the participants before the experiment (according to post-scan task and responses given in run 1). Mediation path coefficients were estimated for each participant independently. Statistical inferences were drawn at the group level for each coefficient using a bias-corrected bootstrap significance test relaxing the normality assumption (10 000 permutations). Averaged paths coefficient and standard deviations are reported directly on Fig. 2D. The algorithm could not converge for 3 participants, which were excluded, due to their high percentage of correct responses (above 90%). The condition (answers repeated or not during the trivia task) and the presence of prior knowledge (watched or unwatched status of the target movie) were included as covariates of non-interest.

*Statistical inferences*

When testing correlation or difference between two variables, parametric statistical tests (i.e Pearson correlation coefficient and Student t-test) were used when the two variables were normally distributed. Their non-parametric equivalent was used otherwise (i.e Spearman rank correlation and Wilcoxon u-test).

### Supplementary FIGURES

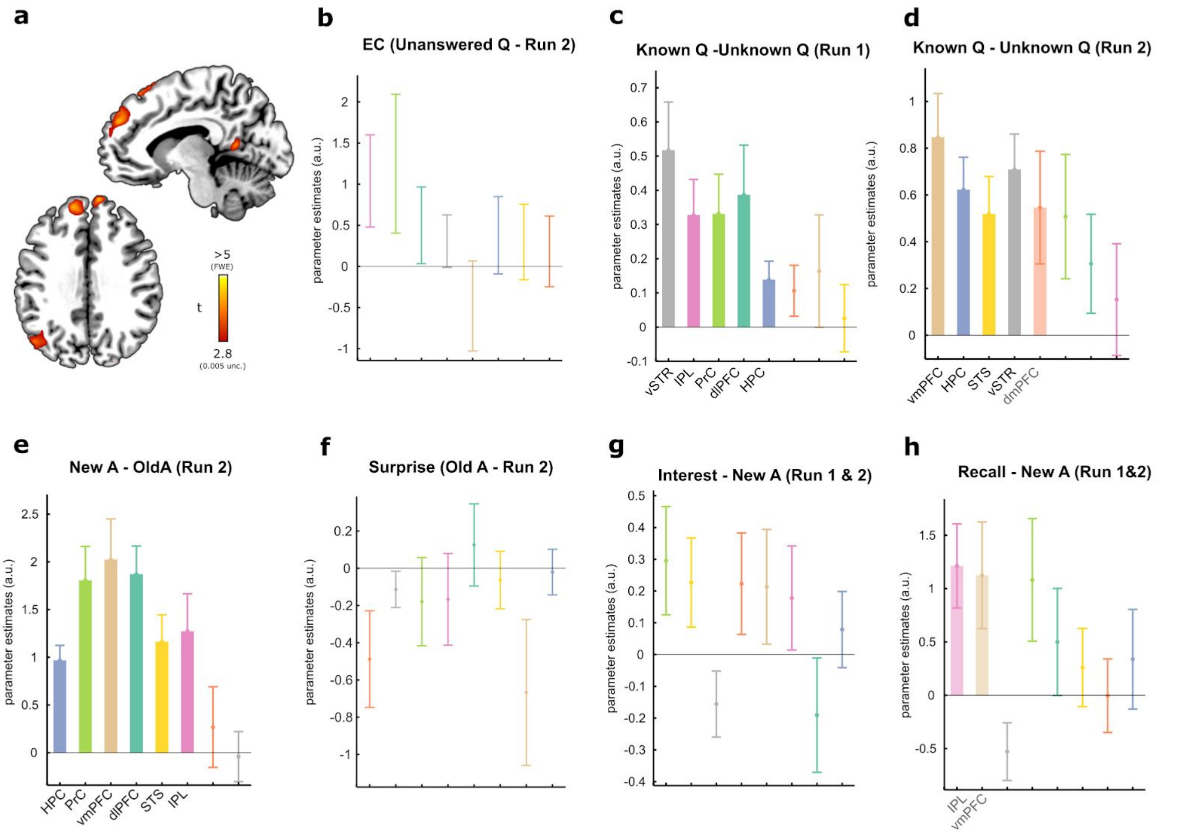


**Supplementary Figure 1.** Additional fMRI results. (a) In the first run, the dmPFC encoded curiosity at the question stage (voxel-wise threshold: p<0.005, cluster-wise threshold: p<0.05FWE). (b) In the second run, none of the ROIs (see Fig. 5b and Methods for a description) was encoded curiosity at the question stage (parameter estimates in the dmPFC are shown in orange). (c) and (d) In both runs, questions who answer were known before the experiment (run 1) or spontaneously recalled from run 1 (run 2) elicited stronger responses in the ventral striatum and the hippocampus. Other ROIs responded to this contrast depending on the context. (e) All the structures which were significant for the contrast “answer > hashtag” in run 1 were also significant for the contrast “New answers > Repeated and remembered answers” in run 2 (compared with Fig. 5a-b). (f) In line with the very notion of surprise, surprise ratings did not correlate with any ROI for repeated and remembered answers in run 2. Due to counterbalancing of items repeated or not across participants, this negative result excludes several confounds related to the visual or linguistic properties of answers (compare with Fig. 5f). (g) Interest ratings did not relate to vmPFC activity (compare with Fig. 5C-F). (h) Items subsequently remembered in the post-scan memory test elicited greater activity in the inferior parietal lobule and in the vmPFC than forgotten items, although the effect did not survive correction for multiple comparison.

For each bar plot, areas surviving the correction for multiple comparisons (across ROIs, pFDR<0.05) are plotted with plain colors, areas significant only at an uncorrected threshold (p<0.05) are plotted with half-transparent colors and non-significant effects are reported using only error bars. Effect are ordered from left to right as a function of their significance. Shaded areas and error bars represent s.e.m.

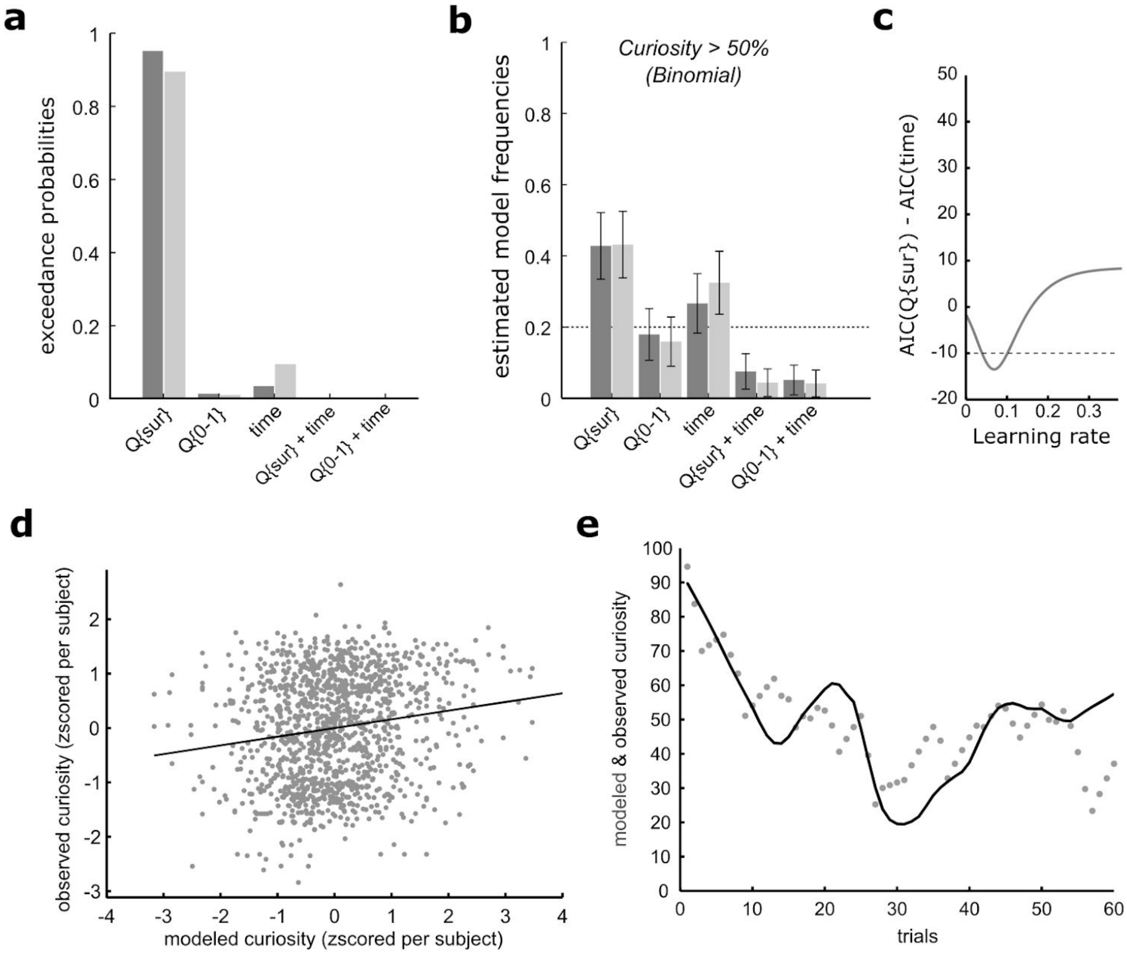


**Supplementary Figure 2.** Additional modelling results. (a) Exceedances probabilities resulting from Bayesian group comparisons favored the model which updated expected surprise on each trial (Q{sur}: AIC-based comparison in dark grey: ep=0.95; BIC in light grey: ep=0.90). The same conclusion held for binarized ratings (AIC: ep=0.82; BIC ep=0.73). (b) Model frequencies estimated when curiosity ratings were binarized (see also Fig. 4a). Note that, when summed over all participants (hence treating model attribution as a fixed effect), AIC and BIC values also strongly favored Q{sur}: as compared to the second most prevalent model ("time only"), both metrics were reduced by more than 10 units for continuous curiosity (BIC difference: 11.1, AIC difference: 13.2) and by more than 6 units for binomial curiosity (BIC difference: 6.8, AIC difference: 8.9), hence reflecting very strong (>10) and strong (>6) evidence for the best-fitting model, respectively (Kass and Raftery, 1995). (c) Advantage of the best-fitting model (Q{sur}) over the second best-fitting alternative (time only) represented as a function of the model learning rate. The best-fitting learning rate (0.069) was consistent with a slowly fluctuating representation of average surprise affecting curiosity levels. (d) Correlation between observed and modeled curiosity ratings pooled over all subjects (r=0.15, p<0.001). Values were z-scored in participant independently in order to exclude irrelevant inter-individual variance. (e) Representation of modeled (grey dots) and observed curiosity levels (black line, smoothed for display purposes) in one participant. Lower values indicate better fit. Error bars correspond to s.e.m.

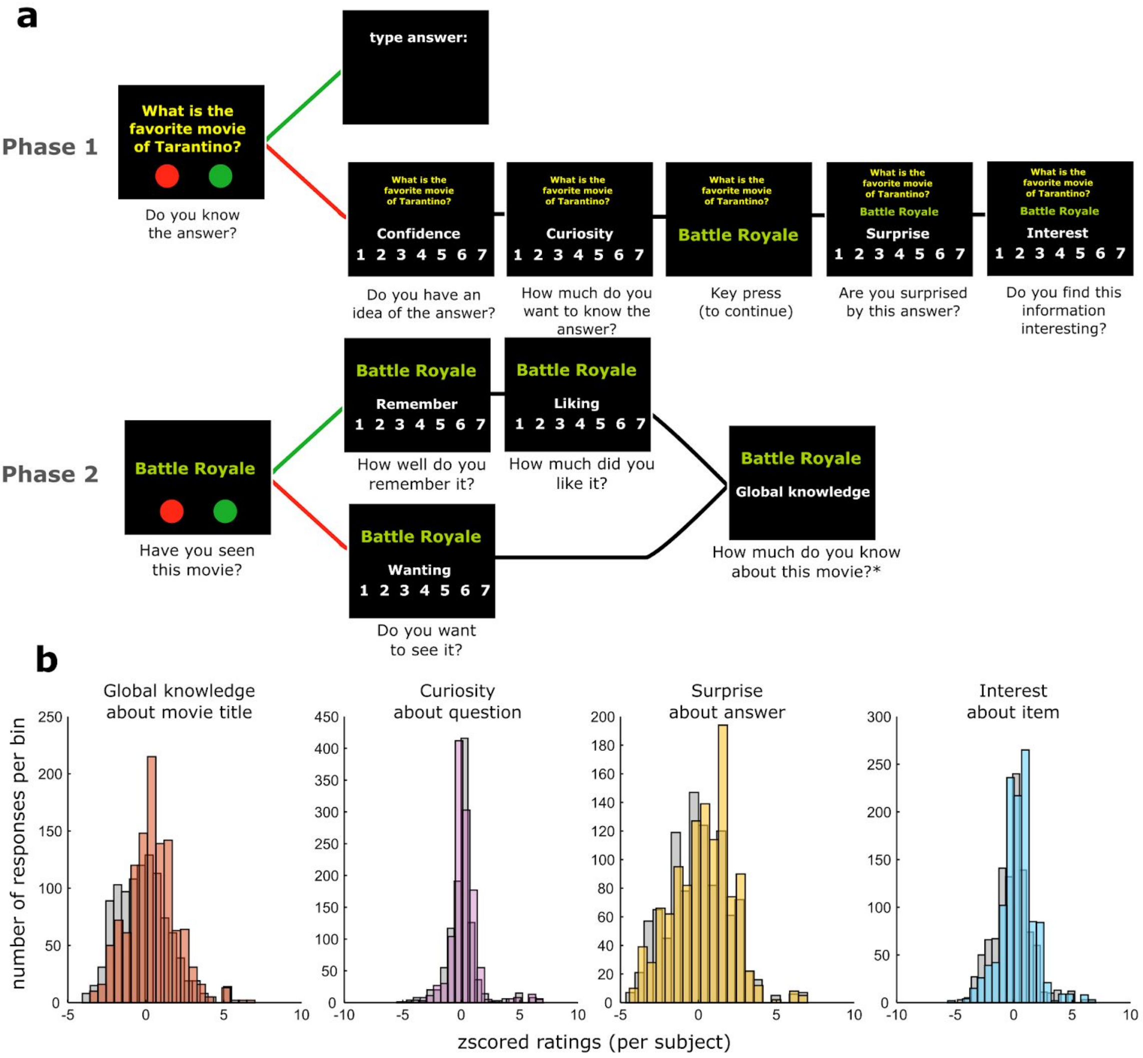


**Supplementary Figure 3.** Behavioural experiment performed to pre-screen trivia items. (a) Sixty-four participants had to evaluate a series of trivia questions (40 per participant) for which the answer was always a movie title. First, participants indicated whether or not they knew the answers (13.2±11.5%). If they knew the answer, they were asked to type it on the keyboard. Otherwise, they had to indicate how confident they were about their best guess, as well as how curious they were to see the answer. Afterwards the answer was displayed and participants rated how surprising they found the answer to be, as well as how interesting it was. In a second step, participants were asked whether or not they had watched each target movie. If they answered yes, they were asked to rate how much they could remember about it and how much they liked it. Otherwise, they rated how much they would like to watch it. Finally, participants reported the amount of knowledge they had about the movie. (b) For inclusion in the fMRI experiment, we selected items to maximise levels prior knowledge, surprise and interest while maintaining a large range of curiosity levels.

| |
|---|
| What movie is presented as a true story in the opening, and as a fiction in the closing credits? |
| From wich movie a crucial scene about crash statistics was cut by most airlines on their trips? |
| What movie was censored by its own director after a series of crimes that were inspired by it? |
| For what movie did Sean Connery refuse a role saying that he did not understand the script? |
| On what movie did Pedro Almodovar work for more than a year before abandoning the project? |
| What movie stands first in the ideal film librairy of the magazine "Les Cahiers du cinéma"? |
| For what movie Italian women gave up to 200kg of hair to make the wigs used by the extras? |
| What movie title is etymologically derived from a word whose meaning is "brotherly love"? |
| What movie what shot as a color movie but was released as a black and white movie? |
| What is the favourite movie of Quentin Tarantino among movies released since 1992? |
| What movie from 1999 required the director to shoot more than 1500 reels of film? |
| What was the first animated movie that was nominated for the Oscar of Best movie? |
| For what movie did Eddie Vedder from Pearl Jam win a Golden Globe for Best Song? |
| In what movie does Clint Eastwood perform the theme song in the closing credits? |
| What hyper violent movie from 1974 was released only in 1999 in British cinemas? |
| In what movie does Michael Keaton improvise most of the lines of his character? |
| For what movie four different endings were shot and submitted to the producers? |
| In what movie from 1994 did Samir Naceri appear for the first time in a movie? |
| What movie from 1983 actually reveals true aspects of Coluche's private life? |
| What movie was released in Quebec with the translated title "Brillantine"? |
| For what movie Forest Whitaker went so far as to learn a new language? |
| In what movie does the director make some winks to the Picsou comics? |
| For what movie Jacques Chirac had a private projection at the Elysée? |
| For what movie of science fiction Will Smith refused the title role? |
| In what movie Uma Thurman interprets the role of the goddess Venus? |
| With what movie Joel Coen won a Palme d'or at the Cannes Festival? |
| What was the first comedy released in Germany after World War II? |
| What movie title was directly inspired by a poem from Edgar Poe? |
| What movie is a free adaptation of Isaac Asimov's novels? |
| What was the favourite movie of the dictator Adolf Hitler? |

| |
|---|
| For what movie did Michelle Pfeiffer turn down the title role because she found the script scary? |
| What movie was shot 15 years after the script was written to benefit from better special effects? |
| What American movie was inspired from Alexandre Dumas' novel "The Lady of the Camellias"? |
| What movie title refers to a hobby consisting in spotting all of a certain type of rolling stock? |
| What movie was critized because its director took some liberties with the sexuality of the hero? |
| For what movie Stanley Kubrick almost received the Razzie Award of worse director of the year? |
| For what movie was Marlon Brando paid four millions dollars for a ten minute performance? |
| In what movie title the question mark doesn't appear because it was considered bad luck? |
| What is the only movie of the James Bond series that was approved by Chinese censorship? |
| For what american movie did Raymon Queneau translate and adapt the dialogues in French? |
| For what movie the director asked that the public shouldn't be allowed to arrive late? |
| For what movie from 1939 did a Black American actress win a Oscar for the first time? |
| For the shooting of what movie Tom Cruise's career was frozen for almost three years? |
| What movie is presented as a postmodern adapatation of George Orwell's novel"1984"? |
| In what action movie did the director sacrifice his own car for budgetary reasons? |
| What movie is a modern adaptation of Jane Austeen's novel Pride and Prejudice? |
| What movie title was inspired by a mispelling made by the son of her director? |
| On what movie did Steven Spielberg refuse to receive any payment for his work? |
| What movie from 1940 was banned in Spain until Francisco Franco died in 1975? |
| What is the first movie from the Cohen brothers that was adapted from a book? |
| For what movie released in 1997 did Jennifer Anniston refuse the title role? |
| In what French movie David Cameron makes a brief apparition of 30 seconds? |
| For what movie did Monica Belluci receive her first role outside of Italy? |
| For what movie Bourvil, who died before the shooting, had to be replaced? |
| What movie was released in Quebec with the translated title "Décadence"? |
| In what movie did Lee Van Cleef appear for the first time in a movie? |
| In what movie Sigourney Weaver appears for the first time on screen? |
| In what movie a real building of Fox Company was partly destroyed? |
| What movie was originally the final year project of its director? |
| In what movie the clocks are blocked at 16h20 in all scenes? |

**Supplementary Table 1 (related to Fig. 1 and S3).** The two lists of trivia questions translated from French. The association between lists and conditions(answer delivered or not in run 1) was counterbalanced across participants.

| | Operationalization | Comment |
|---|---|---|
| **Epistemic curiosity** | **Rating for each item (0 to 100, run 1)** | |
| **Baseline EC** | **Intercept of the model** | **Relates to trait curiosity** |
| **Unspecific EC** | **$\beta_{average\ surprise}$ from the delta-rule algorithm (time-varying, item-independent)** | **Relates to “diversive” or “interest” curiosity** |
| **Specific EC** | **ε ⇔ error of the model (item-dependent)** | **Relates to knowledge gap & “deprivation” curiosity** |
| **Epistemic surprise** | **Rating for each item (1 to 5, post-scan test)** | **Corresponds to unexpected information associated with answers.** |
| **Expected surprise** | **Trial by trial variable derived from the model-based analysis of EC** | **Corresponds to the amount of surprise recently experienced.** |
| **Prior knowledge** | **Target movie watched or not (0/1, based on pre-scan test)** | **Captures episodic knowledge about answers, confounded with prior willful choices.** |
| **Interest** | **Items categorized in the 30 “most interesting” or 30 “least interesting”** | **Captures a global valuation signal about trivia items** |
| **Recall** | **Questions for which answers could be recalled or not in the post-test (0/1)** | **Reflects the efficiency of memory encoding** |
| **Novelty** | **In run 2, contrast between answers never revealed minus answers revealed in run 1 but forgotten.** | **Captures the treatment and encoding of truly novel information.** |
| **Tip of the tongue** | **In run 2, contrast between questions answered but forgotten in run 1 minus questions answered and recalled.** | **Pre-retrieval categorization of potentially known and unknown questions.** |

**Supplementary Table 2.** The two lists of trivia questions translated from French. The association between lists and conditions (answer delivered or not in run 1) was counterbalanced across participants.

### Supplementary references


Ahsan, R.L., Allom, R., Gousias, I.S., Habib, H., Turkheimer, F.E., Free, S., Lemieux, L., Myers, R., Duncan, J.S., Brooks, D.J., Koepp, M.J., Hammers, A., 2007. Volumes, spatial extents and a probabilistic atlas of the human basal ganglia and thalamus. Neuroimage 38, 261–270.

Daunizeau, J., Adam, V., Rigoux, L., 2014. VBA: a probabilistic treatment of nonlinear models for neurobiological and behavioural data. PLoS Comput. Biol. 10, e1003441.

Gläscher, J., 2009. Visualization of group inference data in functional neuroimaging. Neuroinformatics 7, 73–82.

Gruber, M.J., Gelman, B.D., Ranganath, C., 2014. States of curiosity modulate hippocampus-dependent learning via the dopaminergic circuit. Neuron 84, 486–496.

Kang, M.J., Hsu, M., Krajbich, I.M., Loewenstein, G., McClure, S.M., Wang, J.T.-Y., Camerer, C.F., 2009. The wick in the candle of learning: epistemic curiosity activates reward circuitry and enhances memory. Psychol. Sci. 20, 963–973.

Kass, R.E., Raftery, A.E., 1995. Bayes Factors. J. Am. Stat. Assoc. 90, 773–795.

Litman, J.A., Spielberger, C.D., 2003. Measuring epistemic curiosity and its diversive and specific components. J. Pers. Assess. 80, 75–86.
